# The integrated stress response induces R-loops and hinders replication fork progression

**DOI:** 10.1101/2020.03.11.987131

**Authors:** Josephine Ann Mun Yee Choo, Denise Schlösser, Valentina Manzini, Anna Magerhans, Matthias Dobbelstein

**Author notes:** Corresponding author. Correspondence and requests for materials should be addressed to M. D.

## Abstract

The integrated stress response (ISR) allows cells to rapidly shut down most of their protein synthesis in response to protein misfolding, amino acid deficiency, or virus infection. These stresses trigger the phosphorylation of the translation initiation factor eIF2alpha, which prevents the initiation of translation. Here we show that triggering the ISR drastically reduces the progression of DNA replication forks within one hour, thus flanking the shutdown of protein synthesis with immediate inhibition of DNA synthesis. DNA replication is restored by compounds that inhibit eIF2alpha kinases or re-activate eIF2alpha. Mechanistically, the translational shutdown blocks histone synthesis, promoting the formation of DNA:RNA hybrids (R-loops) which interfere with DNA replication. Histone depletion alone induces R-loops and compromises DNA replication. Conversely, histone overexpression or R-loop removal by RNaseH1 each restores DNA replication in the context of ISR and histone depletion. In conclusion, the ISR rapidly stalls DNA synthesis through histone deficiency and R-loop formation. We propose that this shutdown mechanism prevents potentially detrimental DNA replication in the face of cellular stresses.

**SIGNIFICANCE:** The integrated stress response has long been explored regarding its immediate impact on protein synthesis. Translational shutdown represents an indispensable mechanism to prevent the toxicity of misfolded proteins and virus infections. Our results indicate that the shutdown mechanisms reach far beyond translation and immediately interfere with DNA synthesis as well. ISR depletes cells of new histones which induce accumulation of DNA:RNA hybrids. The impairment of DNA replication in this context supports cell survival during stress.

Our work provides a link between the ISR and another subject of active research, i. e. the regulatory network of DNA replication forks.

Graphical Abstract

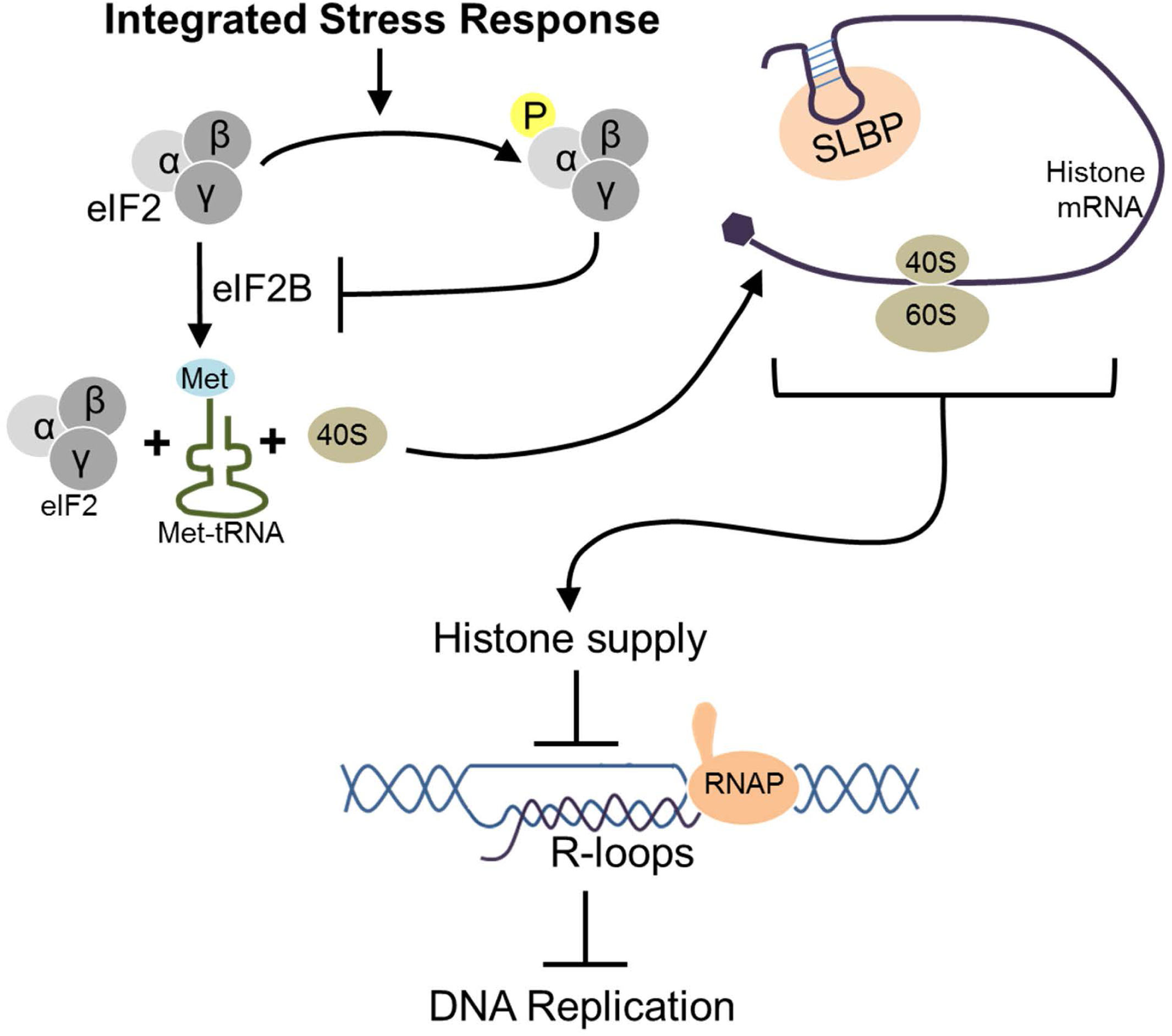

## INTRODUCTION

The integrated stress response (ISR) is widely known as a mechanism to shut down the synthesis of most proteins when the cell suffers various stresses (1) through the activation of the following kinases. Protein kinase R (PKR) is activated upon virus infection and accumulation of double-stranded RNA. PKR-like endoplasmic reticulum kinase (PERK) becomes active when unfolded proteins accumulate in the endoplasmic reticulum. General control nonderepressible 2 (GCN2) responds to amino acid deprivation. And heme-regulated inhibitor (HRI) is triggered in the case of heme depletion in erythrocytes. Each of these kinases triggers the phosphorylation of the alpha subunit of translation initiation factor eIF2 at Serine 51 (2). This modification of eIF2 shuts down the translation of most mRNAs, with the exception of a few mRNAs that employ alternative mechanisms of translation initiation. One of these exceptions is the transcription factor ATF4, which is synthesized with greater efficiency as part of the ISR (3, 4) and then triggers a transcriptional program to counteract the specific stress stimuli (5). The ISR thus prevents further damage to the cell by avoiding further protein synthesis in the context of proteotoxic stress, or as part of a defense mechanism against virus infection or nutrient depletion.

Besides gene expression, the replication of DNA represents an extreme demand on the cell with regard to metabolic activity and energy consumption. For one round of DNA replication, each human cell must synthesize and incorporate 2×3×10^9^ dNTPs. This raises the question whether the ISR might also affect the replication of DNA, perhaps protecting the cell in the context of nutrient deprivation or infection. And indeed, the replication of DNA is a highly regulated process. Regulation is not only implied by the control of cell cycle progression. Rather, even during S phase, the cell can stall the progression of replication forks (6). One example of the underlying mechanisms is provided by the kinase MAPKAPK2, the activation of which diminishes replication fork progression (7, 8). Also, the absence of the tumor suppressor p53 or its target gene product Mdm2 can each enhance replication stress (9, 10). Another way of slowing down DNA replication consists in the lack of histone supply, e.g. by depleting histone chaperones (11, 12). In this situation, the newly synthesized DNA can no longer associate with nucleosomes to a sufficient extent. By mechanisms that are currently not fully explained, this leads to a reduction in DNA synthesis (11–13). Finally, replication stress can be induced by the formation of R-loops, i.e. DNA:RNA hybrids that form by the looping out the non-template strand of DNA after transcription, allowing the newly synthesized RNA to rehybridize with its template strand (14, 15). Such R-loops represent obstacles to DNA replication (16–18).

Previous findings provided hints that the ISR might not only affect the synthesis of proteins but also that of DNA (19, 20), with the earlier report mainly focusing on the drug thapsigargin and its role in replication through interfering with calcium homeostasis. On the other hand, Cabrera *et al*., uses thapsigargin to hinder proper protein folding (induce “ER stress”) which subsequently impaired firing of origins and hence overall DNA synthesis (20). The mechanism was suggested to occur through the activation of claspin and its associated kinase Chk1 (20). Moreover, cycloheximide, a compound that inhibits overall protein synthesis, was found to diminish histone synthesis and slow down DNA replication (12, 21). This raises the question whether the ISR might generally interfere with DNA replication progression, through a shortage of histone synthesis.

Here we show that the ISR triggered by various kinases each interferes with the progression of DNA replication forks. This can be mimicked by the depletion of histones. Strikingly, the removal of R-loops by RNaseH1, or the overexpression of histones, restores DNA replication upon ISR. In addition, histone depletion alone led to an accumulation of R-loops. This suggests a general mechanism that links ISR to the impairment of replication forks, through histone depletion and R-loops.

## MATERIALS & METHODS

### LEAD CONTACT AND MATERIALS AVAILABILITY

Further information and requests for resources and reagents should be directed to and will be fulfilled by the Lead Contact Matthias Dobbelstein.

This study did not generate unique reagents.

### EXPERIMENTAL MODEL AND SUBJECT DETAILS

#### Cell culture

The human osteosarcoma cell line U2OS (p53 proficient, female) was purchased from ATCC (RRID:CVCL_0042). Cells were maintained in Dulbecco’s modified Eagle’s medium (DMEM) supplemented with 10% fetal bovine serum (Merck), 2 mM L-glutamine (Life Technologies), 50 units/ml penicillin, 50 μg/ml streptomycin (Gibco), and 10 µg/ml Ciprofloxacin (Bayer) at 37°C in a humidified atmosphere with 5% CO_2_. Cells used were routinely tested and ensured to be negative for mycoplasma contamination.

## METHOD DETAILS

### Treatments and transfections

Cells were treated with thapsigargin (Thap, Sigma), 1H-Benzimidazole-1-ethanol, 2,3-dihydro-2-imino-alpha-(phenoxymethyl)-3-(phenylmethyl)-monohydrochloride (BEPP, Sigma), L-Histidinol (L-Hist, Sigma), (E)-2-(2-Chlorobenzylidene) hydrazinecarboximidamide (Sephin, Sigma), trans-N,N′-(Cyclohexane-1,4-diyl)bis(2-(4-chlorophenoxy) acetamide (Integrated stress response inhibitor or ISRIB, Sigma), GSK2606414 (PERK inhibitor or PERK i, Calbiochem), gemcitabine (Gem, Actavis), Cycloheximide (CHX, Sigma), 5,6-Dichloro-1-β-D-ribofuranosylbenzimidazole (DRB, Sigma) or LDC067 (Selleckchem) as indicated in the figure legends. Thap, BEPP, Sephin, ISRIB, PERK i, DRB and LDC067 were dissolved in DMSO, L-Hist and gemcitabine dissolved in water, and CHX was dissolved in 100% ethanol.

siRNA transfections were performed using Lipofectamine 3000 (Life Technologies). Cells were reverse transfected with 100 nM siRNA against SLBP (Ambion, custom made, pool of 3 siRNAs) or negative control scrambled siRNA (Ambion, pool of 2 siRNAs), medium replenished after 24 h and cells harvested 40 h post-transfection. For plasmid overexpression, 2 µg of the respective plasmids were forward transfected using Lipofectamine 2000. Medium was replenished after 6 h, and cells were harvested for experiments 24 h post-transfection. The following plasmids were used.

| **Plasmid** | **Origin** |
| --- | --- |
| pICE-NLS-mCherry | Addgene #60364 |
| pICE-RNaseH1-NLS-mCherry | Addgene #60365 |
| pICE-RNaseH1-D10R-E48R-NLS-mCherry | Addgene #60367 |
| pFRT-ToDest-FlagHA | Addgene #26361 |
| pFRT-ToDest-FlagHA-RNaseH1 | Addgene #65782 |
| pCDNA3.1-Flag-H2A | Addgene #63560 |

### Cell synchronization

To obtain a majority population of cells in S phase, cells were synchronized using double thymidine block. Briefly, cells were seeded accordingly and allowed to settle and attach onto plates or coverslips for at least 6 h, then treated with 2 mM thymidine (Sigma). After 16 h, cells were washed once in PBS and then replenished with fresh DMEM for 8 h prior to the second thymidine block (2 mM) for another 16 h. Depending on the assay, cells were released into fresh DMEM for 1 h (celigo proliferation assay) or 4 h (R-loop detection on cells treated with CHX) prior to treatment, harvest and analysis.

### Immunoblot analysis

Cells were washed once in PBS and harvested in radioimmunoprecipitation assay (RIPA) lysis buffer (20 mM TRIS-HCl pH 7.5, 150 mM NaCl, 10 mM EDTA, 1% Triton-X 100, 1% deoxycholate salt, 0.1% SDS, 2 M urea) in the presence of protease inhibitors. Samples were briefly sonicated to disrupt DNA-protein complexes. The protein extracts were quantified using the Pierce BCA Protein assay kit (Thermo Scientific Fisher). Protein samples were boiled at 95°C in Laemmli buffer for 5 min, and equal amounts were analyzed by sodium dodecyl sulfate polyacrylamide gel electrophoresis (SDS-PAGE). Subsequently, proteins were transferred onto a nitrocellulose membrane, blocked in 5% (w/v) non-fat milk in PBS containing 0.1% Tween-20 for 1 h and incubated with primary antibodies at 4°C overnight followed by incubation with peroxidase-conjugated secondary antibodies (donkey anti-rabbit or donkey anti-mouse IgG, Jackson Immunoresearch). The proteins were detected using either Super Signal West Femto Maximum Sensitivity Substrate (Thermo Fisher) or Immobilion Western Substrate (Millipore).

Soluble histones were extracted as described (12). Briefly, cells were washed once in PBS and harvested in a low detergent, hypotonic buffer (10 mM Tris, pH 7.4, 2.5 mM MgCl_2_, and 0.5% NP-40) for 10 min on ice. Following centrifugation at 1000 x *g*, the concentration of the solubilized proteins was determined as described above and equal amounts were analyzed by SDS-PAGE.

| **Antibodies** | **Source (Catalog number)** | **Research Resource Identifiers (RRID)** |
| --- | --- | --- |
| ATF4 (D4B8) | Cell Signaling (#11815) | RRID:AB_2616025 |
| Chk1 | Cell Signaling (#2360) | RRID:AB_2080320 |
| eIF2alpha | Cell Signaling (#9722) | RRID:AB_2230924 |
| Flag | Sigma (F1804) | RRID:AB_262044 |
| gamma H2AX, γH2AX | Cell Signaling (#2577) | RRID:AB_2118010 |
| H3 | Abcam (ab1791) | RRID:AB_302613 |
| H3K56ac | Cell Signaling (#4243) | RRID:AB_10548193 |
| H4K5ac (EP1000Y) | Abcam (ab51997) | RRID:AB_2264109 |
| H4K12ac (EPR17906) | Abcam (ab177793) | RRID:AB_2651187 |
| HSC70 | Santa Cruz (sc-7298) | RRID:AB_627761 |
| mCherry | Abcam (ab167453) | RRID:AB_2571870 |
| phospho-Chk1 (S317) | Cell Signaling (#2344) | RRID:AB_331488 |
| phospho-eIF2alpha (S51) | Cell Signaling (#9721) | RRID:AB_330951 |
| RNaseH1 | Abcam (ab56560) | RRID:AB_945244 |
| SLBP (EPR12673) | Abcam (ab181972) | N/A |

### DNA fiber assay

DNA fiber assays were performed as described previously (9). Briefly, cells were incubated with 5-chloro-2’-deoxyuridine (CldU, Sigma Aldrich) for 30 min, followed by 60 min incubation with 5-iodo-2’-deoxyuridine (IdU, Sigma Aldrich) in the presence of inhibitors or treatments as indicated. For the 7-label assay, cells were incubated with CldU for 1 h and then pulsed labelled with IdU and CldU for 15 min each for a duration of 1.5 h.

Cells were lysed using spreading buffer (200 mM Tris pH 7.4, 50 mM EDTA, 0.5% SDS) and DNA fiber spread on glass slides prior to fixation in a methanol:acetic acid solution (3:1). Upon treatment with 2.5 M HCl, fibers were incubated with rat anti-BrdU antibody (Abcam, RRID:AB_305426, 1:1000, to detect CldU) and mouse anti-BrdU (Becton Dickinson, RRID:AB_10015219, 1:400, to detect IdU) for 1 h at room temperature, then fixed with 4% paraformaldehyde in PBS for 10 min. Slides were incubated with Alexa Fluor 555-conjugated goat anti-rat IgG antibody (RRID:AB_141733) and Alexa Fluor 488-conjugated goat anti-mouse IgG antibody (RRID:AB_138404) (both from Thermo Fisher, 1:200) for 2 h at room temperature.

### S9.6 Immunofluorescence

Cells were seeded on glass coverslips, transfected or treated with reagents accordingly and fixed with 4% paraformaldehyde in PBS for 10 min. Then, cells were permeabilized with 0.5% Triton X-100 in PBS for 15 min, blocked with 3% bovine serum albumin (BSA) in PBS containing 0.1% Tween-20 for 1 h and incubated overnight at 4°C with S9.6 antibody (Kerafast, RRID:AB_2687463, 1:100, to detect DNA:RNA hybrids). Coverslips were washed in PBS prior to incubation with Alexa Fluor 488-conjugated donkey anti-mouse IgG antibody (Thermo Fisher, RRID:AB_141607, 1:250) for 2 h and subsequently counterstained with 0.5 µg/ml DAPI (Sigma) for 5 min prior to mounting using the Fluorescent Mounting Medium from DakoCytomation (#S302380-2) and imaged.

### Dot blot analysis

Cells were seeded, treated with Thap, BEPP or CHX as indicated and harvested. Prior to CHX treatment, cells were synchronized using double thymidine block as described (chapter “Cell synchronization”) and released into fresh DMEM for 4 h prior to addition of CHX. Cells were washed once in PBS and fixed with 1.1% paraformaldehyde in a solution of 0.1 M NaCl, 1 mM EDTA, 0.5 mM EGTA and 50 mM HEPES pH 7 for 30 min at room temperature. To quench the cross-linking reaction, glycin was added to a final concentration of 0.125 M for 5 min. Subsequently, the cells were lysed in 1% Triton X-100, 0.15 M NaCl, 1 mM EDTA, 0.3% SDS with protease inhibitors. The cell lysates were sonicated for 10 cycles (30 sec on/off) (Bioruptor, Diagenode) and then subjected to 2 mg/ml proteinase K (Thermo Fisher) treatment for 1 h at 50°C. DNA was isolated using phenol-chloroform extraction and DNA concentration normalized between samples.

The DNA (1.3 µl) was spotted onto pre-wet nitrocellulose membrane, allowed to air dry and then cross-linked with UVC for 5 min. The membrane was blocked in 5% BSA in PBS containing 0.25% Tween-20 for 30 min at room temperature and subsequently incubated with S9.6 antibody (Kerafast, 1:300) in blocking solution overnight at 4°C. Following incubation with peroxidase-conjugated donkey anti-mouse IgG (Jackson Immunoresearch, RRID:AB_2340773,1:10000), DNA:RNA hybrids (as measured using S9.6 intensity) were detected using Super Signal West Femto Maximum Sensitivity Substrate (Thermo Fisher). To confirm the specificity of the antibody, one half of the DNA samples were also pre-treated with RNaseH (0.03 U/ng DNA, Ambion Thermo Fisher) for 3 h at 37°C prior to spotting. As a loading control, the membrane was subsequently incubated with antibodies to single-stranded DNA (ssDNA). Briefly, the membrane was incubated with 2.5 M HCl for 15 min (to denature the DNA), washed with PBS, and incubated with antibody to ssDNA (Millipore, RRID:AB_570342, 1:1000) for 2 h at room temperature. The detection of ssDNA was performed following exposure to secondary antibody using Super Signal West Femto Maximum Sensitivity Substrate (Thermo Fisher).

### EdU Incorporation Assay

5-ethynyl-2’-deoxyuridine (EdU, Thermo Fisher Scientific, #A10044) was added to exponentially growing cells to a final concentration of 20 µM for 1 h until harvest. Prior to imaging, the cells were fixed and permeabilized as done for immunofluorescence staining. The following reagents were added to 100 mM Na-Phosphate buffer (pH 7) in the following order: 5 µM Alexa Fluor 488 picolyl-azide or 5 µM Alexa Fluor 594 picolyl-azide (Jena Biosciences, #CLK-1276-1 or #CLK-1296-1), 100 µM CuSO_4_ (Jena Biosciences, #CLK-MI004) in 500 µM tris-hydroxypropyltriazolylmethylamine (THPTA; Sigma-Aldrich, #762342) and 5 mM Na-Ascorbate (Jena Biosciences, #CLK-MI005). The click reaction was performed for 1 h on a shaker, at room temperature and protected from light. Samples were subsequently washed thrice for 10 min with PBS, followed by incubation with 0.3 µg/ml DAPI (Sigma-Aldrich, #D9542) for 10 min. For experiments in Fig. 1, cells were kept in PBS prior to image acquisition with the Celigo Imaging Cytometer (Nexcelom Bioscience). DAPI was used to create a nuclear mask and quantify the DNA content while the nuclear EdU signal was quantified using the Celigo image analysis software. EdU and DAPI signals were presented in a horseshoe plot. For experiments in Fig. 4, cover slips were mounted using the fluorescent mounting medium (DakoCytomation, #S302380-2) and imaged.

**FIGURE 1:**
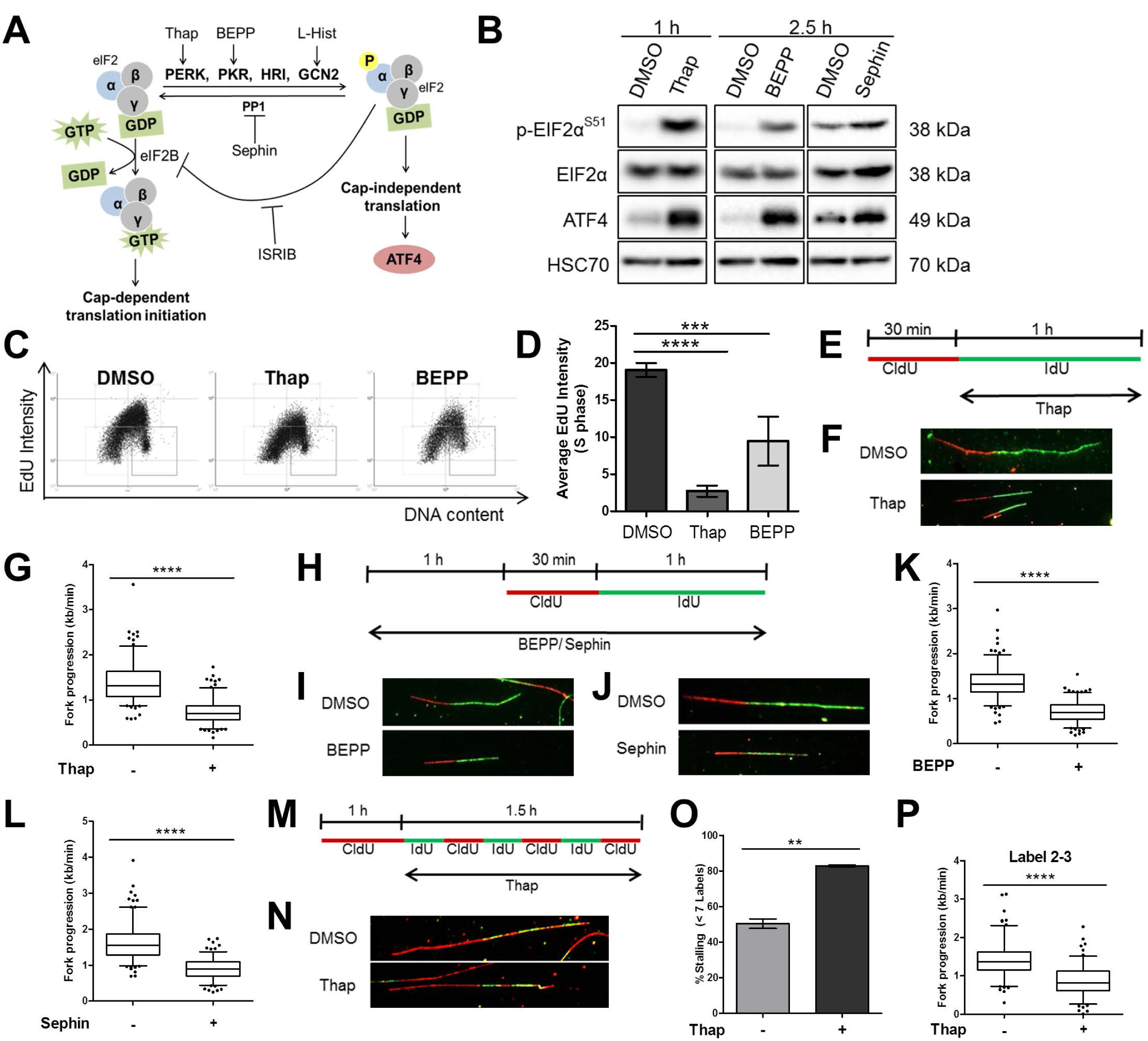
DNA replication is compromised shortly after ISR induction. **(A)** Schematic representation of the ISR that can be activated upon stimulation of the kinases PERK, PKR or GCN2 or upon inhibition of the phosphatase PP1 using thapsigargin, BEPP-monohydrochloride, L-Histidinol or Sephin respectively. Activation of ISR can be measured by an increase in eIF2alpha phosphorylation or by the accumulation of ATF4. ISR can be inhibited using a small molecule inhibitor, ISRIB. **(B)** Immunoblot analysis of cells treated with Thap (4 μM), BEPP (10 μM) or Sephin (25 μM) to confirm ISR induction. HSC70 as loading control. **(C)** Representative horseshoe plots showing EdU incorporation in relation to DNA content (DAPI) of cells treated with DMSO, Thap (4 μM, 1 h) or BEPP (10 μM, 2.5 h). **(D)** Average EdU staining intensity of cells in S phase as determined from the plots in (**C**) and displayed as mean ± SD. For second replicate, see Fig. S1B. **(E)** U2OS cells were incubated with 5’-chloro-2’-deoxy-uridine (25 μM CldU, 30 min) followed by 5-iodo-2’-deoxyuridine (250 μM IdU, 60 min) in the presence of 4 μM Thap prior to harvesting for DNA fiber analysis. **(F)** Representative labeled tracks of newly synthesized DNA incorporating CldU (red) and IdU (green) of cells in (**E**). **(G)** Fork progression as determined from IdU track length (kb/min), displayed as 5-95 percentile whiskers boxplot of Thap–treated cells. Box plots represent data from one out of 3 independent experiments. See Fig. S1C,D for additional experiments. **(H)** U2OS cells were pre-treated with 10 μM BEPP or 25 μM Sephin for 1 h and subsequently incubated with CldU (30 min) and IdU (60 min) in the presence of these reagents and then harvested for analysis. **(I/J)** Representative fiber tracks as visualized by immunostaining of CldU (red) and IdU (green) of BEPP (**I**) or Sephin (**J**) −treated cells. **(K/L)** Fork progression calculated from the IdU label (kb/min) of BEPP (**K**) or Sephin (**L**) –treated cells. Fork progression displayed as boxplots with 5-95 percentile whiskers which are representative of one out 3 independent experiments. See Fig. S1E–H. **(M)** Cells were pulsed labeled with CldU (25 μM, 60 min) and then alternately with IdU (25 μM) and CldU (25 μM) for 15 min intervals for a duration of 1.5 h in the presence of Thap (4 μM), then harvested for 7-label fiber assay analysis (9). From this, the number of labels incorporated was used for fork stalling analysis and the length of labels 2-3 was used for fork progression analysis. **(N)** Representative images of fiber tracks that have incorporated 7 labels. **(O)** Percentage of forks with less than 7 labels indicating higher fork stalling rate of cells treated with Thap. Chart represents mean ± SD of two independent experiments. **(P)** Velocity of fork determined from track length of labels 2 to 3 displayed as box plots (5-95 percentile whiskers). Plot is a representative of 2 independent experiments. See Fig. S1M.

### Proliferation assay (Celigo)

To study the long-term effect of ISR on cells in S phase, proliferation assay was conducted on synchronized cells. Cells were seeded in technical duplicates in 24-well plates, synchronized using double thymidine block (as described), and released into fresh medium for 1 h then treated with BEPP (30 µM) for 6 h to ensure ISR activation during S phase of the cells. During synchronization, cells were also transfected with plasmids to RNaseH1 or an empty vector control as described previously. After 6 h of treatment, medium was replenished and confluency of cells at day 0 was measured using Celigo Imaging Cytometer (Nexcelom Bioscience). Measurements were made subsequently every 24 or 48 h and medium was changed prior to every measurement.

### QUANTIFICATION AND STATISTICAL ANALYSIS

#### DNA fiber analysis

To avoid bias, data acquisition and analysis were conducted in a double-blinded manner where identities of the samples were blinded prior to imaging and analysis. Whenever possible, a minimum of 100 DNA fiber structures were visualized with fluorescence microscopy (Axio Scope A1 microscope (Zeiss) equipped with an Axio Cam MRc/503 camera) and analyzed.

For the 7-label fiber assay, the number of labels incorporated was counted using the cell counter plugin on Fiji. Fork stalling was then calculated by dividing the number of tracks with less than all seven labels by the total number of tracks and converted into percentage. The length of the second to third label was measured to determine the replication progression for the 7-label fiber assay. The Fiji software (RRID:SCR_002285) (22) was used to measure the labeled tracks in pixels and converted to micrometers using the conversion factor of 1 µm = 5.7 pixels (as determined by measuring scale bar under the same microscope settings) and then to kilo base (kb) using the conversion factor 1 µm = 2.59 kb. Rate of fork progression was calculated by dividing the number of bases by the labeling time of the track.

For the 2-label fiber assays, fibers were analyzed for their IdU track length and IdU fork progression rate calculated as described.

#### Nuclear quantification of immunofluorescence

Images were acquired (same exposure time for all images for each fluorescent channel per experiment) with Axio Scope A1 microscope (Zeiss) equipped with an Axio Cam MRc/503 camera.

The Fiji software was used for automated analysis and quantification of nuclear S9.6 or EdU staining. DAPI staining was used to identify regions of interest (nuclei) prior to measuring mean intensity of the Alexa Fluor 488 staining (S9.6), Alexa Flour 488 picolyl-azide or Alexa Fluor 594 picolyl-azide (EdU). At least 200 cells were subjected to analysis and quantification.

#### Statistical testing

Statistical testing was performed using Graph Pad Prism 6 (RRID:SCR_002798). For fiber assay and immunofluorescence experiments, a Mann–Whitney U test was used to calculate significance. For the other experiments, an unpaired Student’s t-test was calculated. Significance was assumed where p-values ≤ 0.05. Asterisks represent significance in the following way: ****, p ≤ 0.0001, ***, p ≤ 0.005; **, p ≤ 0.01; *, p ≤ 0.05.

## RESULTS

### DNA replication is compromised shortly after ISR induction

The ISR triggers a shutdown of protein synthesis, representing an emergency response to nutrient deprivation or proteotoxic stress. Here, we tested whether this response might also affect the synthesis of DNA. We induced the ISR and the consequent phosphorylation of eIF2alpha at Serine 51 by stimulating the kinases PERK, PKR and GCN2, or by inhibiting GADD45A (regulatory subunit of the PP1 phosphatase) using the small compounds thapsigargin (Thap) (23), BEPP-monohydrochloride (24), L-Histidinol (25) or Sephin (26) respectively (Fig. 1A). Increased phosphorylation of eIF2alpha and elevated expression of ATF4 following treatment confirmed ISR activation in all cases (Fig. 1B; Fig. S1A). Sephin inhibits the removal of constitutive phosphate modifications on eIF2alpha. This induces a moderate increase in phosphorylation of eIF2alpha, less pronounced than with Thap or BEPP, i.e. activators of eIF2 alpha kinases. We first performed an EdU incorporation assay to measure overall DNA synthesis in individual cells upon ISR activation during S phase. As shown (Fig. 1C,D; Fig. S1B), the activation of ISR using Thap or BEPP significantly reduced DNA synthesis in S phase. Then, we measured the progression of single DNA replication forks using DNA fiber assays, measuring the length of DNA tracks with incorporated IdU (Fig. 1E). Treatment with Thap led to a reduction in fork progression (Fig. 1F,G; Fig. S1C,D). In addition, we found that treatment of U2OS cells with BEPP, Sephin or L-Histidinol all impaired DNA fork progression significantly, albeit to different extents (Fig. 1H–L; Fig. S1E–L). To understand if the reduction in fork progression upon ISR was due to lower speed of DNA polymerase or a higher frequency of polymerase stalling, we conducted a 7-label fiber assay on Thap−treated cells (Fig. 1M) as described in our previous publications (9, 10). This revealed both increased stalling of DNA polymerase (i.e. decreased processivity) and slower DNA polymerization (Fig. 1N−P; Fig. S1M).

Interestingly, despite the significant reduction in DNA replication progression following ISR stimulation, we did not observe a substantial increase in phosphorylation of Chk1 or histone variant H2AX (gamma H2AX) (Fig. S1N) as compared to gemcitabine, a well-established inducer of replicative stress (7) indicating that the ISR slows down replication forks without triggering a strong DNA damage response. These results suggest that the ISR not only triggers a shutdown in protein synthesis but also imposes severe and immediate restrictions on DNA replication.

### Pharmacological antagonists of ISR partially rescue DNA replication

Based on our findings suggesting that the ISR interferes with DNA replication, we now investigated if these effects are downstream of phosphorylated eIF2alpha and could be reversed using a small molecule inhibitor of ISR known as ISRIB (27–29) (Fig. 1A). ISRIB enhances the activity of the nucleotide exchange factor eIF2B, thereby overcoming the inhibitory effect of eIF2alpha phosphorylation. We pre-treated cells with ISRIB, followed by the ISR inducers Thap, BEPP or Sephin, and then measured DNA replication fork progression (Fig. 2A,B). Single treatment of cells with Thap, BEPP or Sephin resulted in an impairment of DNA replication as observed before, but pre-treatment of these cells with ISRIB significantly prevented this inhibition of DNA replication (Fig. 2C–H; Fig. S2A−E). Similarly, inhibition of PERK with a pharmacological inhibitor, PERKi or GSK2606414 (30), was also able to significantly rescue DNA replication defects by Thap treatment (Fig. S2F−J). Activation and inhibition of ISR were confirmed using ATF4 detection as readout (Fig. S2K,L). These findings clarify that the compounds used interfere with DNA replication through the ISR and through eIF2alpha phosphorylation.

**FIGURE 2:**
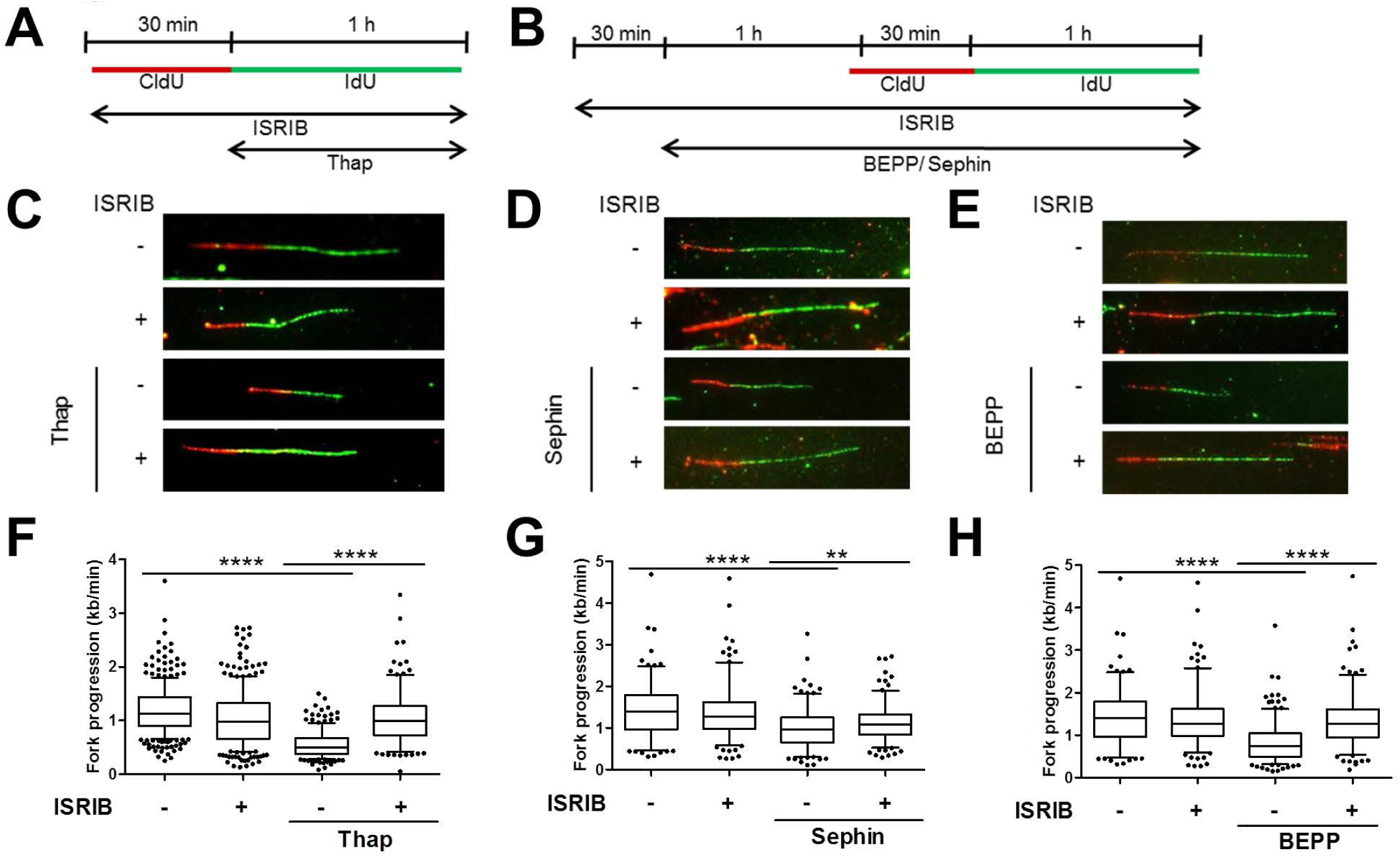
Pharmacological antagonists of ISR partially rescue DNA replication. **(A)** U2OS cells were treated with 1 μM ISRIB and at the same time incubated with CldU (30 min). Cells were labeled with IdU (60 min) in the presence of ISRIB and 4 μM Thap and then harvested for DNA fiber assay analysis. **(B)** Cells were pre-treated with 1μM ISRIB for 30 min and then with 10 μM BEPP or 25 μM Sephin in the presence of ISRIB for 2.5 h. To label newly synthesized DNA, cells were incubated with CldU and IdU during the last 1.5 h as shown, and then harvested for analysis. **(C−E)** Representative DNA tracks as labeled in red (CldU) and green (IdU) of cells treated with ISRIB/Thap (**C**), ISRIB/Sephin (**D**), or ISRIB/BEPP (**E**). **(F−H)** Fork progression of IdU label of cells treated with ISRIB/Thap (**F**), ISRIB/Sephin (**G**) or ISRIB/BEPP (**H**) represented as 5-95 percentile box plots. Plots shown are a representative of 2 or 3 independent experiments. See Fig. S2.A–E.

### Stimulation of the ISR induces R-loops

We were now searching for a mechanism that allows the ISR to interfere with DNA replication. DNA:RNA hybrids (R-loops) have recently emerged as one of the major players in regulating DNA replication (15, 17, 31). They are formed through the hybridization of newly-synthesized RNA to its template DNA while looping out the opposite DNA strand. R-loops can pose as a steric hindrance to an oncoming replisome, thereby blocking DNA replication (16). We investigated if ISR induction led to an enrichment of R-loops. In cells treated with Thap or BEPP, we detected DNA:RNA hybrids by immunofluorescence with an antibody against them (S9.6) (Fig. S3A) (32, 33). As a negative control, we overexpressed RNaseH1 (33) in these cells, i. e. an RNase that specifically removes the RNA portion of DNA:RNA hybrids (Fig. S3A) (15). By quantification, we found a significant increase in the intensity of S9.6 fluorescence in the nuclei of cells treated with Thap or BEPP (Fig. 3A; Fig. S3B). Upon RNaseH1 overexpression, the S9.6 staining intensity within these nuclei decreased to intensities similar to control−treated cells (Fig. 3A; Fig. S3B). We confirmed RNaseH1 overexpression and ISR induction by immunoblot analysis of RNaseH1 and ATF4 levels (Fig. S3C). To supplement our immunofluorescence experiments, we performed dot blot analyses using the antibody S9.6. Cells were treated with Thap or BEPP, followed by chromatin preparation. Samples were also treated with RNaseH as a negative control. In each case, DNA:RNA hybrids were then detected on dot blots. Similar to the immunofluorescence, we observed a strong increase in S9.6 intensity upon ISR, which was abolished by RNaseH (Fig. 3B,C; Fig. S3D,E). Thus, ISR activation leads to an enrichment of R-loops.

**FIGURE 3:**
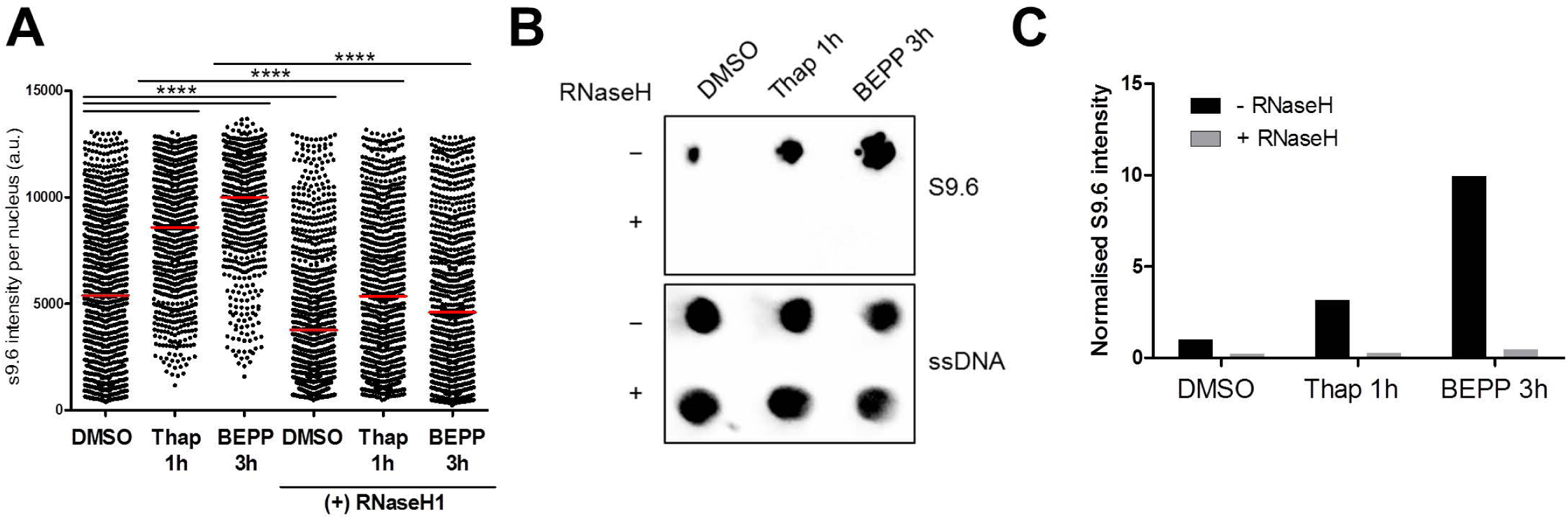
Stimulation of the ISR induces R-loops. **(A)** Cells were treated with 4 μM Thap or 10 μM BEPP for 1 h or 3 h respectively with/without RNaseH1 overexpression prior to fixation and S9.6 immunofluorescence analysis. Scatter plot of S9.6 intensity per nucleus of cells (arbitrary units), determined by quantification from one of 2 independent experiments (see Fig. S3A,B). Red line represents mean nuclear S9.6 staining. **(B)** ISR was induced in cells using Thap (4 μM, 1 h) or BEPP (10 μM, 3 h), followed by dot blot analysis to quantify DNA:RNA hybrids. Equal amounts of DNA were spotted onto nitrocellulose membrane, and R-loops were detected using the S9.6 antibody. RNaseH treatment was conducted alongside and used as a negative control to confirm specificity of the signal. The signal of ssDNA was used as an internal sample loading control. See Fig. S3D for a replicate. **(C)** The S9.6 signals obtained in (**B**) were quantified, normalized against the loading control (ssDNA signal), then against DMSO (without RNaseH) and plotted as bar charts. See Fig. S3E.

### Removal of R-loops re-establishes DNA replication upon induction of ISR but compromises survival of stressed cells

As ISR activation induced more R-loops, we hypothesized that these R-loops were responsible for compromising DNA replication. To test this, we first overexpressed wildtype or catalytically mutant RNaseH1 (32) and treated these cells with Thap or BEPP to induce ISR, and measured total DNA synthesis by EdU incorporation. As seen previously (Fig. 1C,D), EdU incorporation was reduced upon ISR (Fig. 4A; Fig. S4A−C). Strikingly, we now observed that the overexpression of catalytically active RNaseH1 largely restored EdU incorporation and thus DNA synthesis in both Thap and BEPP−treated cells (Fig. 4A; Fig. S4A−C). To test if removal of R-loops was also able to rescue single DNA fork progression, we subjected cells overexpressing wildtype or catalytically inactive RNaseH1and treated with Thap or BEPP to DNA fiber assay analysis (Fig. 4B; Fig. S4F). The removal of R-loops with wildtype but not mutant RNaseH1 completely rescued DNA replication in the context of ISR (Fig. 4C,D; Fig. S4D−J). Immunoblot analysis confirmed that RNaseH1 overexpression did not interfere with eIF2alpha phosphorylation (Fig. S4K,L) and thus not with the ISR *per se*. We then hypothesized that R-loop induction and the resulting impairment of DNA replication upon ISR might help cells to survive by halting the complex DNA replication program in the face of stress conditions. To investigate if the inhibition of DNA replication following accumulation of R-loops upon ISR is protective to the cell, we conducted a proliferation assay of cells treated with BEPP in the presence or absence of RNaseH1. Indeed, removal of R-loops *via* the overexpression of RNaseH1 further reduced proliferation of cells compared to cells that were treated with BEPP alone (Fig. 4E; Fig. S4M,N). Our findings therefore suggest that ISR impairs DNA replication through inducing R-loops and that this inhibition in DNA replication is supporting cell survival during stress.

**FIGURE 4:**
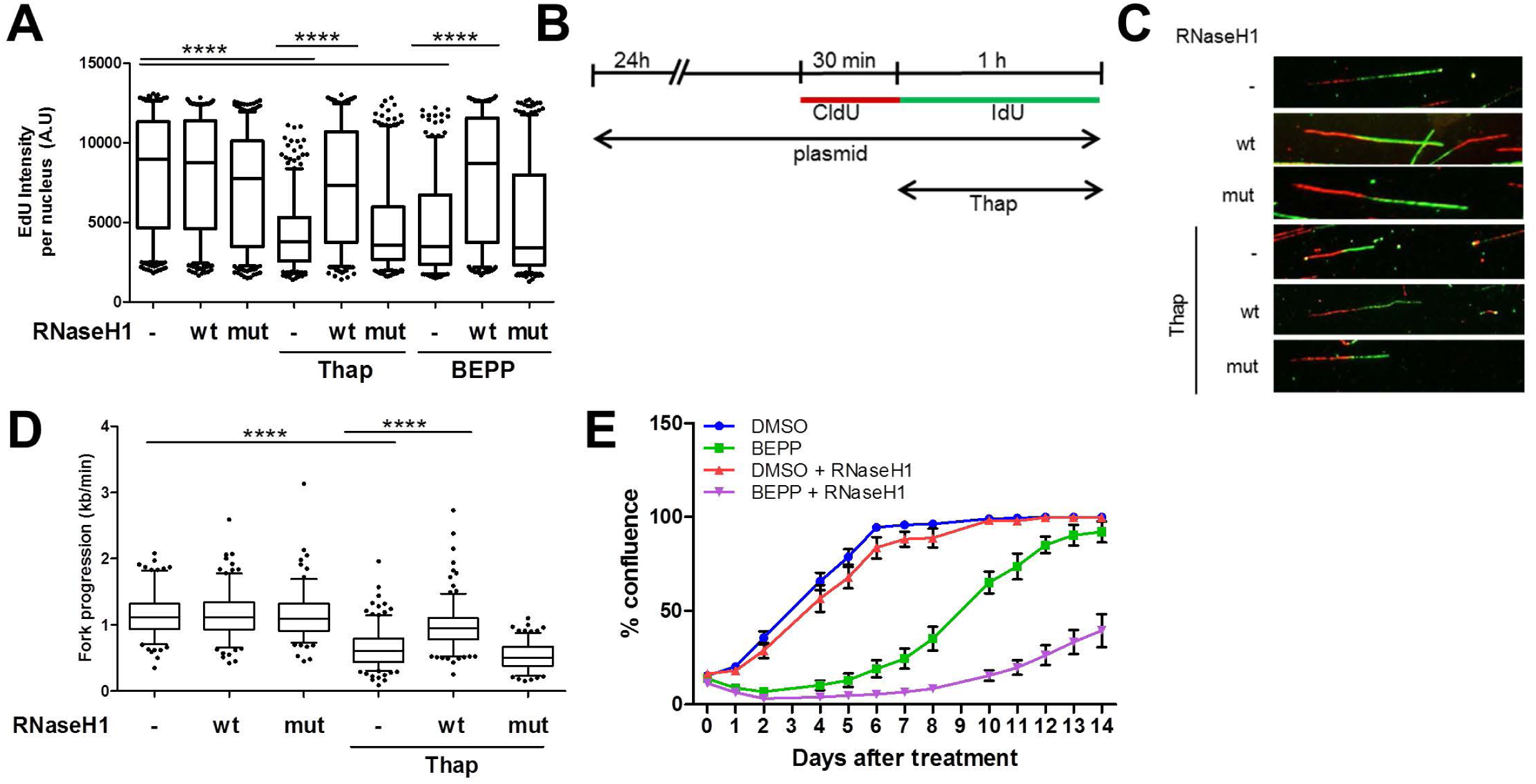
Removal of R-loops re-establishes DNA replication upon induction of ISR but compromises survival of stressed cells. **(A)** U2OS cells transfected with control, RNaseH1 wildtype (wt) or catalytically mutant RNaseH1 (D10R-E48R) (mut) expression plasmids for 24 h were treated with Thap (4 μM, 1 h) or BEPP (10 μM, 2.5 h). During the last 1 h of treatment, the cells were labeled with 20 μM EdU and then subjected to fluorescence analysis of EdU incorporation. Box plot (5-95 percentile whiskers) of the quantified EdU intensities per nucleus of one of 3 independent experiments. For replicates, see Fig. S4B,C. **(B)** Transfection of cells with control or RNaseH1 plasmids (wt or mut) were conducted as described in (**A**) 24 h prior to labeling with CldU (30 min) and IdU (60 min). Cells were treated with 4 μM Thap during the IdU label and then harvested for analysis. **(C)** Representative DNA fiber tracks stained for CldU (red) and IdU (green) of Thap–treated cells overexpressing the respective plasmids as described in (**B**). **(D)** Box plot (5-95 percentile whiskers) of DNA fork progression of cells overexpressing RNaseH1 wt or mut plasmids in the presence/absence of Thap. Fork progression was measured using the IdU label and the plot shown is a representative of one out of 3 independent experiments. See Fig. S4D,E. **(E)** Long term proliferation assay of BEPP−treated cells with/without RNaseH1 overexpression displayed as percentage confluence. Transfected cells that were synchronized at S phase were treated with either DMSO or BEPP (30 μM) for 6 h. The media was then replenished and cell confluency at day 0 was measured using the Celigo Cytometer. Confluency was measured on the indicated days for 2 weeks. Mean ± SD of technical duplicates were plotted. Plot is a representation of 3 biological repeats (Fig. S4M,N).

### Ongoing transcription is required for compromising DNA replication by the ISR

R-loops were suggested to form between RNA and its DNA template, shortly after transcription (15). This raised the hypothesis that short-term inhibition of transcription should re-activate DNA synthesis in the context of the ISR. To test this, we employed two different CDK9 inhibitors, DRB (34) and LDC067 (35). CDK9 inhibition is an established way to interfere with the elongation of transcription (36). We measured DNA replication of cells treated with Thap or BEPP, in the presence or absence of CDK9 inhibitors (Fig. 5A,B). And indeed, the inhibition of transcription significantly rescued DNA replication from its impairment by ISR (Fig. 5C−F; Fig. S5A−D), suggesting that ongoing transcription and R-loops formed by ISR are responsible for impairing DNA replication.

**FIGURE 5:**
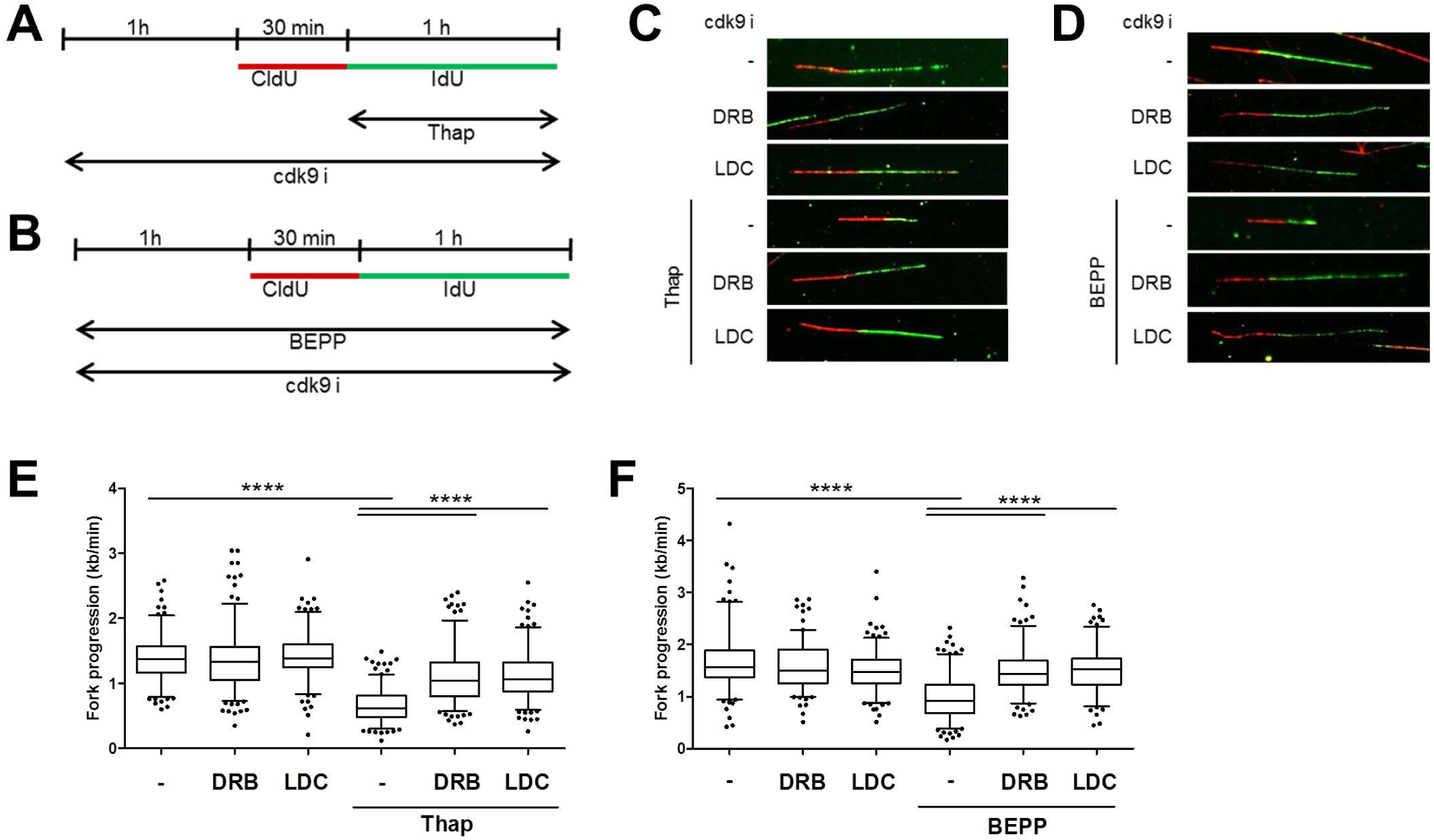
Ongoing transcription is required for compromising DNA replication by the ISR. **(A)** Cells were pre-treated with CDK9 inhibitors (25 μM DRB, 10 μM LDC067) for 1 h prior to labeling with CldU (30 min) and IdU (60 min) in the presence of CDK9i and Thap (4 μM). Cells were then harvested for DNA fiber analysis. **(B)** U2OS cells were treated with CDK9i (25 μM DRB, 10 μM LDC067) and BEPP (10 μM) for 1 h and then labelled with CldU (30 min) and IdU (60 min) with both CDK9i and BEPP prior to analysis. **(C/D**) Representative DNA fiber tracks of cells treated with CDK9i/Thap (**C**) or CDK9i/BEPP (**D**) visualized *via* immunostaining of CldU (red) and IdU (green). **(E/F)** IdU tracks of cells treated with CDK9i and Thap (**E**) or CDK9i and BEPP (**F**) were used to measure fork progression and presented as box plots (5-95 percentile whiskers). One representative plot from 3 independent experiments shown. See Fig. S5A–D.

### ISR activation blocks the synthesis of histones required for DNA replication

Phosphorylation of eIF2alpha at Ser51 during ISR inhibits cap-dependent translation, thereby blocking the synthesis of most proteins in the cell. To investigate if abolished protein synthesis is sufficient to impair DNA replication, we treated cells with a well-established ribosome inhibitor, cycloheximide (CHX), and measured DNA replication progression (Fig. S6A). Within an hour of CHX treatment, we observed a strong reduction in DNA replication (Fig. S6B−D), mimicking the effects we observed with the ISR inducers (Fig 1E−L; Fig. S1C−L). Next, we asked which kind of proteins need to be synthesized continuously to sustain DNA replication. Based on previous reports (11, 12, 21, 37) we suspected that histones need to be provided throughout DNA synthesis to avoid replication stress. Indeed, inducing the ISR by Thap or BEPP quickly reduced the levels of newly synthesized soluble histones, as marked by acetylation of lysine residue 56 on Histone 3 (H3K56ac) or lysine residues 5 or 12 on Histone 4 (H4K5ac or H4K12ac) (12, 38–41), to a similar extent as upon CHX treatment (Fig. 6A; Fig. S6E). To test if a reduction in histone synthesis alone is sufficient to hinder DNA replication in our system as found earlier (12), we used siRNA to deplete the stem loop-binding protein (SLBP) that is required for translation of histones. As expected, SLBP depletion also resulted in a mark decrease in soluble H3K56ac, H4K5ac and H4K12ac (Fig. 6A; Fig. S6E). Of note, a significant impairment in DNA replication was observed by SLBP depletion alone (Fig. S6F−I), strongly suggesting that histones are the critical protein species the reduced synthesis of which is responsible for impaired DNA replication during ISR.

**FIGURE 6:**
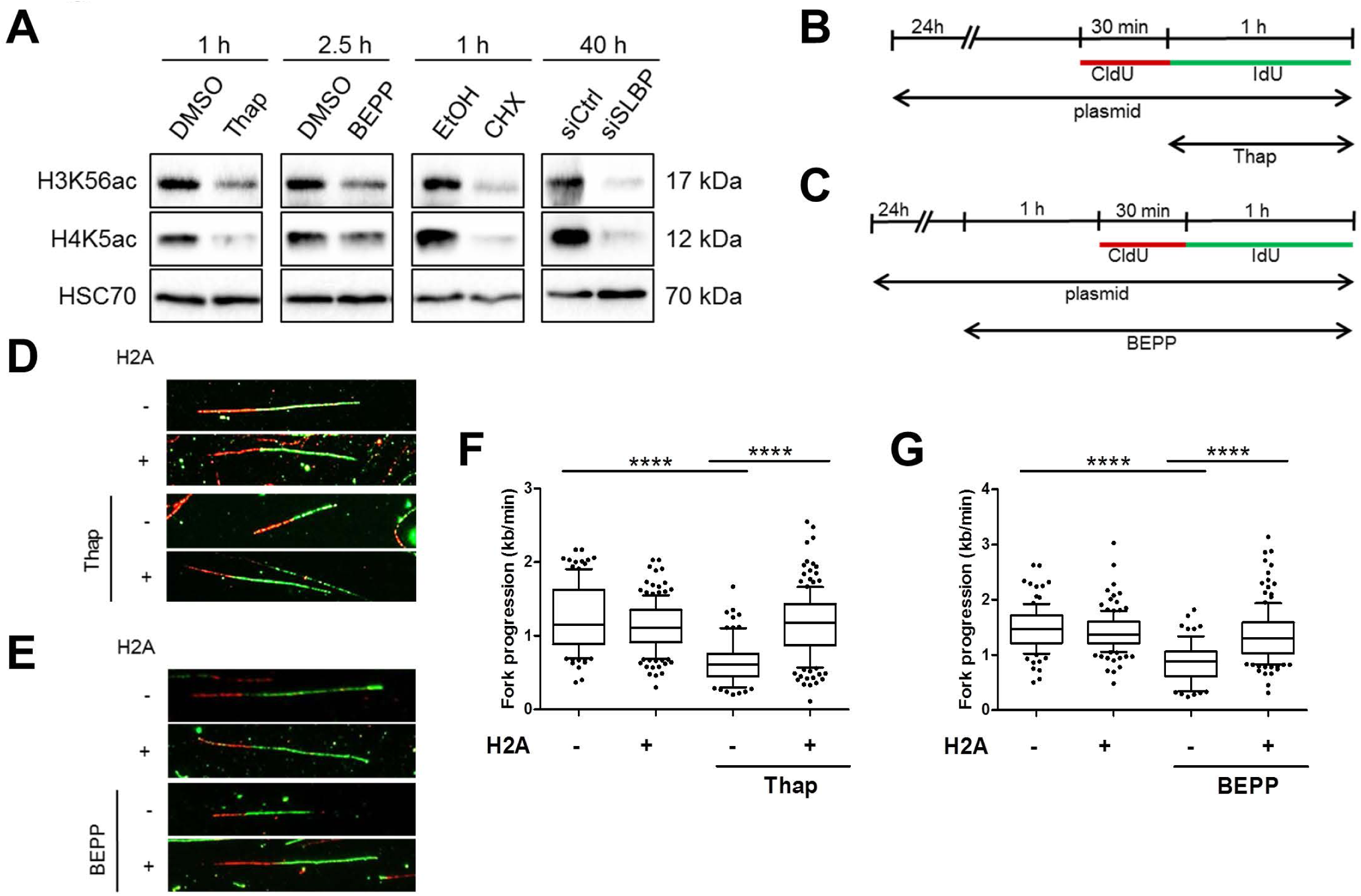
ISR activation blocks the synthesis of histones required for DNA replication. **(A)** Soluble proteins were extracted from cells treated with Thap (4 µM), BEPP (10 µM), CHX (50 µg/ml) or cells transfected with siRNA against SLBP (100 nM). Immunoblot analyses of soluble histone-3 lysine-56 acetylation (H3K56ac) and histone-4 lysine-5 acetylation (H4K5ac) were used to measure newly synthesized histones (12, 39). HSC70 as loading control. **(B)** Cells were transfected with H2A or control plasmids, labeled with CldU (30 min) followed by IdU (60 min). Cells were treated with Thap (4 μM) during the IdU label as indicated prior to analysis. **(C)** Cells were transfected with H2A or empty vector plasmids and treated with BEPP for 2.5 h. Newly synthesized DNA was labeled with CldU (25 μM, 30 min) followed by IdU (250 μM, 60 min) during the last 1.5 h in the presence of BEPP then harvested for analysis. **(D)** Representative DNA fiber tracks of cells transfected with plasmids (control, H2A) and labeled as described in (**B**). **(E)** Images of DNA fibers (representative) of BEPP–treated cells overexpressing control or H2A plasmids visualized as CldU (red) and IdU (green). **(F)** Fork progression (kb/min) of cells in (**D**) calculated using IdU track length. DNA fork progression displayed as box plot (5-95 percentile whiskers) and is a representative data of one of 3 independent experiments. See Fig. S6J,K. **(G)** DNA fork progression (kb/min) of cells treated as in (**C**) and displayed as box plots (5-95 percentile whiskers). IdU label was used to calculate fork progression. Data is a representative of 3 independent experiments. See Fig. S6L,M.

To find out whether restoring histone levels alone might allow DNA replication even during ISR, we measured DNA replication in cells overexpressing histone H2A and treated with Thap or BEPP (Fig. 6B,C). Strikingly, overexpression of histone H2A restored DNA replication despite ISR activation (Fig. 6D−G; Fig. S6J−O). We conclude that the ISR interferes with DNA replication through inhibiting histone synthesis.

### Inhibition of histone synthesis induces R-loops which impairs DNA replication

We have found that the ISR blocks histone synthesis, which compromises DNA replication (Fig. 6). Moreover, the ISR can induce R-loops (Fig. 3), which are also required to perturb DNA replication (Fig. 4; 5). Therefore, we hypothesized that histone deprivation induces the formation of R-loops which then compromises DNA replication. To investigate this, we performed immunofluorescence staining using the S9.6 antibody to detect R-loops on cells that had been treated with CHX to deplete newly synthesized histones, with and without RNaseH1 overexpression. Indeed, CHX−treated cells accumulated DNA:RNA hybrids (Fig. 7A; Fig. S7A−C). Similarly, dot blot analysis using the S9.6 antibody on chromatin from these cells also revealed a profound induction of R-loops which was removed upon RNaseH treatment (Fig. 7B,C; Fig. S7D,E). Next, to investigate if DNA replication impairment by histone depletion could also be restored by removing R-loops, we depleted cells of new histones using CHX or by siRNA to SLBP, in the presence or absence of either wildtype or catalytically inactive RNaseH1, and then measured the progression of DNA replication (Fig. 7D,E). We observed that overexpression of wildtype RNaseH1 but not its mutant rescued DNA replication upon histone depletion (Fig. 7F−I; Fig. S7F−K). Together, these results suggest a mechanistic concept of ISR-induced DNA replication impairment. Accordingly, ISR blocks histone synthesis which then interferes with DNA replication through R-loops.

**FIGURE 7:**
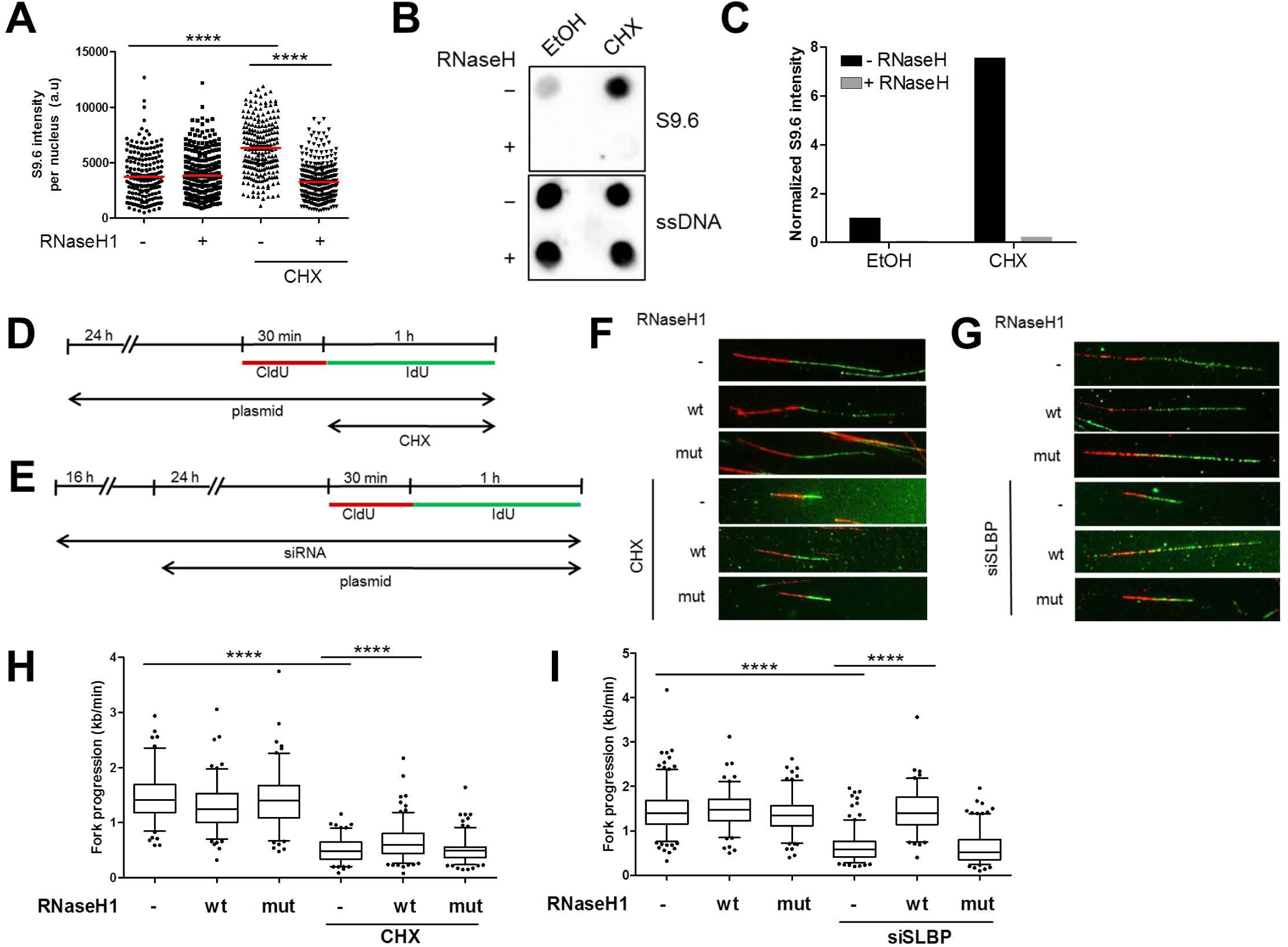
Inhibition of histone synthesis induces R-loops which impairs DNA replication. **(A)** Cells synchronized at S phase and transfected with either control or RNaseH1 expression plasmids were treated with CHX (50 µg/ml) for 1 h, then harvested for S9.6 immunofluorescence analysis as described in Fig. 3A. Intensity of S9.6 staining per nucleus was quantified and displayed as a scatter plot. Red line represents mean S9.6 intensity per nucleus. See Fig. S7B for a replicate. **(B)** Dot blot analysis of S phase cells to detect R-loops using S9.6 antibody. Synchronized cells in S phase were treated with CHX (50 μg/ml, 1 h) and harvested. Equal amounts of DNA were spotted onto nitrocellulose membrane. R-loops were detected using S9.6 antibody whereas subsequent ssDNA detection (on denatured DNA) was used as an internal loading control. As a negative control, samples were treated with RNaseH enzyme for 3 h at 37°C. Blot is a representative of 2 independent experiments. See Fig. S7D. **(C)** Signal from the spots in (**B**) were quantified and normalized to the loading control (ssDNA) and then to the sample without ISR or RNaseH treatment. See Fig. S7E. **(D)** U2OS cells were transfected with plasmids (control, RNaseH1 wt or RNaseH1 mut) 24 h prior to labeling with CldU (30 min) and IdU (60 min). CHX (50 μg/ml) was added to the cell during the IdU label. **(E)** Cells were transfected with siRNA (siCtrl or siSLBP, 100 nM) 16 h prior to overexpression with plasmids (control, RNaseH1 wt or RNaseH1 mut). Cells were then incubated with CldU (30 min) and IdU (60 min) to label newly synthesized DNA and then harvested. **(F/G)** DNA fiber tracks of CHX–treated cells (**F**) or cells depleted from SLBP (**G**) with overexpression of either the control, RNaseH1 wt or RNaseH1 mutant plasmids. Fiber tracks were observed by immunostaining of CldU (red) and IdU (green). **(H/I)** Box plot (5-95 percentile whiskers) showing the fork progression as measured using IdU track length of CHX−treated (**H**) or SLBP–depleted (**I**) cells in the presence/absence of RNaseH1 overexpression. Representative data shown from one of 3 independent experiments. See Fig. S7F−I.

## DISCUSSION

Our results indicate that the ISR compromises DNA replication, within the first hour of eIF2alpha phosphorylation, and through the depletion of histones. When new histones become unavailable, by ISR or histone chaperone inhibition, R-loops mediate the impairment of DNA replication fork progression.

Is this replication stress? Previous reports suggest that the depletion of histones slow down replication fork progression, but do not detectably trigger the activation of Chk1, a classical hallmark of replication stress (6, 12, 21). Similarly, in our hands, Chk1 phosphorylation or phosphorylation of the histone variant H2AX (gamma H2AX) are observed only to a low extent (when compared to treatment with the nucleoside analogue gemcitabine) (Fig. S1N). Taken together with the observed accumulation of R-loops, we conclude that R-loops as such do not necessarily activate Chk1, despite interfering with the progression of DNA replication forks, at least not within the first few hours of blocking DNA replication.

It was previously reported that the lack of histone supply hinders replication fork progression (11, 12, 21). The mechanism(s) were suggested to include interactions of histones with the MCM helicase and/or the delayed removal of PCNA from Okazaki fragments but remain to be fully clarified (12). Our results provide the following explanation. We hypothesize that when histones are missing, nucleosome-free DNA accumulates. This provides more opportunities of DNA:RNA hybridization (Fig. 7A−C). The resulting R-loops turned out to be required for the observed replication fork impairment, since RNaseH1 enhanced DNA synthesis in the context of histone depletion (Fig. 7D−I). However, it remains to be determined how exactly such R-loops lead to stalled replication. Apart from physical collisions, the accumulation of R-loops might trigger signaling pathways that attenuate fork progression (18). Indeed, it has been shown that R-loops induce the phosphorylation of Histone H3 at Ser10 (H3S10), a mark of chromatin compaction (42). It is thus possible that the R-loops formed could lead to torsional stress throughout the DNA surrounding them through chromatin condensation, which then signals the replication machinery ahead to stop replicating DNA (16).

We propose that the inhibition of DNA replication as part of the ISR provides an advantage for cell survival. Under conditions of nutrient deprivation, it is conceivably advantageous that protein synthesis is reduced to a minimum. On top of this, our results show that slowing down DNA synthesis through R-loop accumulation, as a newly established part of the ISR, helps the cell to survive nutrient restriction. This can be seen with a substantial impairment in proliferation of cells overexpressing RNaseH1 under ISR stimulation (Fig. 4E). After all, replicating a diploid human genome within one cell requires 2×3×10^9^ deoxynucleoside-triphosphates, each of which contains two energy-rich anhydride bonds. Stalling replication forks reduces the rate by that dNTPs are used and might thus contribute to survival under conditions of limited available energy. This might have contributed to the evolution of a tight coupling mechanism that immediately shuts down DNA synthesis in the context of ISR.

The ISR has also been suggested as a target for cancer therapy (43, 44). The idea is mainly to exacerbate proteotoxicity and the accumulation of unfolded proteins in cancer cells by inhibitors of kinases that would otherwise stimulate the ISR. Based on the results presented here, it is possible that interfering with the ISR may also overcome the stalling in DNA replication, perhaps enhancing the vulnerability of cancer cells towards drugs that provoke replication stress, e.g. nucleoside analogues or ATR inhibitors (6). This suggests the use of ISR inhibitors with nucleoside analogues and/or ATR inhibitors in an attempt to achieve synergistic responses to eliminate cancer cells.

Proteasome inhibitors and HSP90 inhibitors form part of a general strategy to eliminate cancer cells by targeting essential cellular machineries (45), exploiting non-oncogene addiction (46, 47). However, these inhibitors can induce the ISR as well (48). The results presented here suggest that this will also halt DNA replication forks. It remains to be determined whether this will diminish the activity of DNA-damaging chemotherapeutics towards cancer cells. In such a case, the simultaneous administration of proteotoxic drugs with certain conventional chemotherapeutics might need to be avoided to prevent drug antagonisms. On the other hand, the addition of an ISR inhibitor might restore the cooperation of a proteotoxic and a DNA-damaging drug.

In contrast to the direction explored here, replication stress can also induce the ISR, as has been reported in the case of the nucleoside analogue gemcitabine (49). Of note, however, gemcitabine was found to induce eIF2alpha phosphorylation with a delay of at least 6 hours. In accordance with this, we were also unable to detect eIF2alpha phosphorylation within shorter periods of time upon gemcitabine treatment. Thus, the ISR probably does not affect the immediate response of cells towards direct triggers of replication stress. However, upon long-term application of chemotherapy, the ISR might represent a mechanism of cell resistance, not only by avoiding proteotoxic stress but also by slowing down DNA replication.

Another important aspect of the ISR consists in the defense against virus infection, in particular through activation of the kinase PKR (50, 51). Most obviously, this will reduce the production of virus proteins, e.g. for building new virus particles. Our results suggest that, in addition, DNA synthesis is diminished. On top of cellular DNA, this may also pertain to viral genomes, especially when they are associated with nucleosomes and thus require histone synthesis. This packaging of viral DNA into nucleosomes has been observed (52–54). It is therefore tempting to speculate that the ISR might also contribute to a decrease in the synthesis of viral DNA, perhaps antagonizing virus production more efficiently than through translational shutdown alone.

## ACKNOWLEDGEMENTS

pICE-NLS-mCherry, pICE-RNaseH1-WT-NLS-mCherry and pICE-RNaseH1-D10R-E48R-NLS-mCherry were gifts from Patrick Calsou (Addgene #60364, #60365, #60367). pFRT-ToDest-FlagHA and pFRT-ToDest-FlagHA-RNaseH1 were gifts from Thomas Tuschl (Addgene #26361, #65782). pCDNA3.1-Flag-H2A was a gift from Titia Sixma (Addgene #63560). Our work was supported by the Else Kröner-Fresenius-Stiftung, the Deutsche Krebshilfe, the Wilhelm Sander-Stiftung, the Deutsche José Carreras Stiftung, and the Deutsche Forschungsgemeinschaft. VM was a member of the IMPRS/MSc./PhD program Molecular Biology, and JC was a member of the Göttingen Graduate School GGNB during this work.

## AUTHOR CONTRIBUTIONS

JC and MD conceived the project and designed experiments. JC, DS and AM performed experiments. VM optimized the EdU click reaction. MD and JC wrote the manuscript. All authors read and approved the manuscript.

## REFERENCES

1. Pakos-Zebrucka K, Koryga I, Mnich K, Ljujic M, Samali A, Gorman AM. The integrated stress response. EMBO Rep. 2016;17(10):1374–95.

2. Taniuchi S, Miyake M, Tsugawa K, Oyadomari M, Oyadomari S. Integrated stress response of vertebrates is regulated by four eIF2 alpha kinases. Sci Rep-Uk. 2016;6.

3. Vattem KM, Wek RC. Reinitiation involving upstream ORFs regulates ATF4 mRNA translation in mammalian cells. Proc Natl Acad Sci U S A. 2004;101(31):11269–74. Epub 2004/07/28.

4. Hinnebusch AG. Gene-specific translational control of the yeast GCN4 gene by phosphorylation of eukaryotic initiation factor 2. Mol Microbiol. 1993;10(2):215–23. Epub 1993/10/01.

5. Harding HP, Zhang Y, Zeng H, Novoa I, Lu PD, Calfon M, et al. An integrated stress response regulates amino acid metabolism and resistance to oxidative stress. Mol Cell. 2003;11(3):619–33. Epub 2003/04/02.

6. Dobbelstein M, Sorensen CS. Exploiting replicative stress to treat cancer. Nat Rev Drug Discov. 2015;14(6):405–23.

7. Kopper F, Bierwirth C, Schon M, Kunze M, Elvers I, Kranz D, et al. Damage-induced DNA replication stalling relies on MAPK-activated protein kinase 2 activity. Proc Natl Acad Sci U S A. 2013;110(42):16856–61.

8. Kopper F, Binkowski AM, Bierwirth C, Dobbelstein M. The MAPK-activated protein kinase 2 mediates gemcitabine sensitivity in pancreatic cancer cells. Cell Cycle. 2014;13(6):884–9.

9. Klusmann I, Rodewald S, Muller L, Friedrich M, Wienken M, Li Y, et al. p53 Activity Results in DNA Replication Fork Processivity. Cell Rep. 2016;17(7):1845–57.

10. Klusmann I, Wohlberedt K, Magerhans A, Teloni F, Korbel JO, Altmeyer M, et al. Chromatin modifiers Mdm2 and RNF2 prevent RNA:DNA hybrids that impair DNA replication. Proceedings of the National Academy of Sciences. 2018;115:E11311–E20.

11. Groth A, Corpet A, Cook AJ, Roche D, Bartek J, Lukas J, et al. Regulation of replication fork progression through histone supply and demand. Science. 2007;318(5858):1928–31.

12. Mejlvang J, Feng Y, Alabert C, Neelsen KJ, Jasencakova Z, Zhao X, et al. New histone supply regulates replication fork speed and PCNA unloading. J Cell Biol. 2014;204(1):29–43.

13. Jasencakova Z, Scharf AN, Ask K, Corpet A, Imhof A, Almouzni G, et al. Replication stress interferes with histone recycling and predeposition marking of new histones. Mol Cell. 2010;37(5):736–43.

14. Aguilera A, Garcia-Muse T. R loops: from transcription byproducts to threats to genome stability. Mol Cell. 2012;46(2):115–24. Epub 2012/05/01.

15. Skourti-Stathaki K, Proudfoot NJ. A double-edged sword: R loops as threats to genome integrity and powerful regulators of gene expression. Genes Dev. 2014;28(13):1384–96. Epub 2014/07/06.

16. Santos-Pereira JM, Aguilera A. R loops: new modulators of genome dynamics and function. Nat Rev Genet. 2015;16(10):583–97.

17. Crossley MP, Bocek M, Cimprich KA. R-Loops as Cellular Regulators and Genomic Threats. Mol Cell. 2019;73(3):398–411. Epub 2019/02/09.

18. Garcia-Muse T, Aguilera A. Transcription-replication conflicts: how they occur and how they are resolved. Nat Rev Mol Cell Biol. 2016;17(9):553–63. Epub 2016/07/21.

19. Shukla N, Jeremy JY, Nicholl P, Krijgsman B, Stansby G, Hamilton G. Short-term exposure to low concentrations of thapsigargin inhibits replication of cultured human vascular smooth muscle cells. The British journal of surgery. 1997;84(3):325–30. Epub 1997/03/01.

20. Cabrera E, Hernandez-Perez S, Koundrioukoff S, Debatisse M, Kim D, Smolka MB, et al. PERK inhibits DNA replication during the Unfolded Protein Response via Claspin and Chk1. Oncogene. 2017;36(5):678–86.

21. Henriksson S, Groth P, Gustafsson N, Helleday T. Distinct mechanistic responses to replication fork stalling induced by either nucleotide or protein deprivation. Cell Cycle. 2018;17(5):568–79.

22. Schindelin J, Arganda-Carreras I, Frise E, Kaynig V, Longair M, Pietzsch T, et al. Fiji: an open-source platform for biological-image analysis. Nature methods. 2012;9(7):676–82. Epub 2012/06/30.

23. Thastrup O, Dawson AP, Scharff O, Foder B, Cullen PJ, Drobak BK, et al. Thapsigargin, a novel molecular probe for studying intracellular calcium release and storage. Agents Actions. 1989;27(1-2):17–23. Epub 1989/04/01.

24. Hu W, Hofstetter W, Wei X, Guo W, Zhou Y, Pataer A, et al. Double-stranded RNA-dependent protein kinase-dependent apoptosis induction by a novel small compound. J Pharmacol Exp Ther. 2009;328(3):866–72. Epub 2008/12/11.

25. Hansen BS, Vaughan MH, Wang L. Reversible inhibition by histidinol of protein synthesis in human cells at the activation of histidine. J Biol Chem. 1972;247(12):3854–7. Epub 1972/06/25.

26. Das I, Krzyzosiak A, Schneider K, Wrabetz L, D’Antonio M, Barry N, et al. Preventing proteostasis diseases by selective inhibition of a phosphatase regulatory subunit. Science. 2015;348(6231):239–42. Epub 2015/04/11.

27. Sidrauski C, Acosta-Alvear D, Khoutorsky A, Vedantham P, Hearn BR, Li H, et al. Pharmacological brake-release of mRNA translation enhances cognitive memory. Elife. 2013;2:e00498. Epub 2013/06/07.

28. Tsai JC, Miller-Vedam LE, Anand AA, Jaishankar P, Nguyen HC, Renslo AR, et al. Structure of the nucleotide exchange factor eIF2B reveals mechanism of memory-enhancing molecule. Science. 2018;359(6383). Epub 2018/03/31.

29. Zyryanova AF, Weis F, Faille A, Alard AA, Crespillo-Casado A, Sekine Y, et al. Binding of ISRIB reveals a regulatory site in the nucleotide exchange factor eIF2B. Science. 2018;359(6383):1533–6. Epub 2018/03/31.

30. Axten JM, Medina JR, Feng Y, Shu A, Romeril SP, Grant SW, et al. Discovery of 7-methyl-5-(1-{[3-(trifluoromethyl)phenyl]acetyl}-2,3-dihydro-1H-indol-5-yl)-7H-pyrrolo[2,3-d]pyrimidin-4-amine (GSK2606414), a potent and selective first-in-class inhibitor of protein kinase R (PKR)-like endoplasmic reticulum kinase (PERK). J Med Chem. 2012;55(16):7193–207. Epub 2012/07/26.

31. Gan W, Guan Z, Liu J, Gui T, Shen K, Manley JL, et al. R-loop-mediated genomic instability is caused by impairment of replication fork progression. Genes Dev. 2011;25(19):2041–56. Epub 2011/10/08.

32. Britton S, Dernoncourt E, Delteil C, Froment C, Schiltz O, Salles B, et al. DNA damage triggers SAF-A and RNA biogenesis factors exclusion from chromatin coupled to R-loops removal. Nucleic acids research. 2014;42(14):9047–62. Epub 2014/07/18.

33. Spitzer JI, Ugras S, Runge S, Decarolis P, Antonescu C, Tuschl T, et al. mRNA and protein levels of FUS, EWSR1, and TAF15 are upregulated in liposarcoma. Genes, chromosomes & cancer. 2011;50(5):338–47. Epub 2011/02/24.

34. Baumli S, Endicott JA, Johnson LN. Halogen bonds form the basis for selective P-TEFb inhibition by DRB. Chem Biol. 2010;17(9):931–6. Epub 2010/09/21.

35. Albert TK, Rigault C, Eickhoff J, Baumgart K, Antrecht C, Klebl B, et al. Characterization of molecular and cellular functions of the cyclin-dependent kinase CDK9 using a novel specific inhibitor. Br J Pharmacol. 2014;171(1):55–68. Epub 2013/10/10.

36. Morales F, Giordano A. Overview of CDK9 as a target in cancer research. Cell Cycle. 2016;15(4):519–27. Epub 2016/01/15.

37. Gunjan A, Paik J, Verreault A. Regulation of histone synthesis and nucleosome assembly. Biochimie. 2005;87(7):625–35. Epub 2005/07/02.

38. Alabert C, Groth A. Chromatin replication and epigenome maintenance. Nat Rev Mol Cell Biol. 2012;13(3):153–67. Epub 2012/02/24.

39. Masumoto H, Hawke D, Kobayashi R, Verreault A. A role for cell-cycle-regulated histone H3 lysine 56 acetylation in the DNA damage response. Nature. 2005;436(7048):294–8. Epub 2005/07/15.

40. Groth A, Rocha W, Verreault A, Almouzni G. Chromatin challenges during DNA replication and repair. Cell. 2007;128(4):721–33. Epub 2007/02/27.

41. Sobel RE, Cook RG, Perry CA, Annunziato AT, Allis CD. Conservation of deposition-related acetylation sites in newly synthesized histones H3 and H4. Proc Natl Acad Sci U S A. 1995;92(4):1237–41. Epub 1995/02/14.

42. Castellano-Pozo M, Santos-Pereira JM, Rondon AG, Barroso S, Andujar E, Perez-Alegre M, et al. R loops are linked to histone H3 S10 phosphorylation and chromatin condensation. Mol Cell. 2013;52(4):583–90. Epub 2013/11/12.

43. Ojha R, Amaravadi RK. Targeting the unfolded protein response in cancer. Pharmacol Res. 2017;120:258–66.

44. Urra H, Dufey E, Avril T, Chevet E, Hetz C. Endoplasmic Reticulum Stress and the Hallmarks of Cancer. Trends Cancer. 2016;2(5):252–62.

45. Dobbelstein M, Moll U. Targeting tumour-supportive cellular machineries in anticancer drug development. Nat Rev Drug Discov. 2014;13(3):179–96.

46. Luo J, Solimini NL, Elledge SJ. Principles of cancer therapy: oncogene and non-oncogene addiction. Cell. 2009;136(5):823–37.

47. Nagel R, Semenova EA, Berns A. Drugging the addict: non-oncogene addiction as a target for cancer therapy. EMBO Rep. 2016;17(11):1516–31.

48. Suh DH, Kim MK, Kim HS, Chung HH, Song YS. Unfolded protein response to autophagy as a promising druggable target for anticancer therapy. Ann N Y Acad Sci. 2012;1271:20–32.

49. Palam LR, Gore J, Craven KE, Wilson JL, Korc M. Integrated stress response is critical for gemcitabine resistance in pancreatic ductal adenocarcinoma. Cell Death Dis. 2015;6:e1913.

50. Balachandran S, Roberts PC, Brown LE, Truong H, Pattnaik AK, Archer DR, et al. Essential role for the dsRNA-dependent protein kinase PKR in innate immunity to viral infection. Immunity. 2000;13(1):129–41. Epub 2000/08/10.

51. Garcia MA, Gil J, Ventoso I, Guerra S, Domingo E, Rivas C, et al. Impact of protein kinase PKR in cell biology: from antiviral to antiproliferative action. Microbiol Mol Biol Rev. 2006;70(4):1032–60. Epub 2006/12/13.

52. Knipe DM, Lieberman PM, Jung JU, McBride AA, Morris KV, Ott M, et al. Snapshots: chromatin control of viral infection. Virology. 2013;435(1):141–56.

53. Bock CT, Schranz P, Schroder CH, Zentgraf H. Hepatitis B virus genome is organized into nucleosomes in the nucleus of the infected cell. Virus Genes. 1994;8(3):215–29. Epub 1994/07/01.

54. Lieberman PM. Chromatin organization and virus gene expression. J Cell Physiol. 2008;216(2):295–302. Epub 2008/03/05.

