## Supplementary figures for "The integrated stress response induces R-loops and hinders replication fork progression"

### Supplementary Figure S1

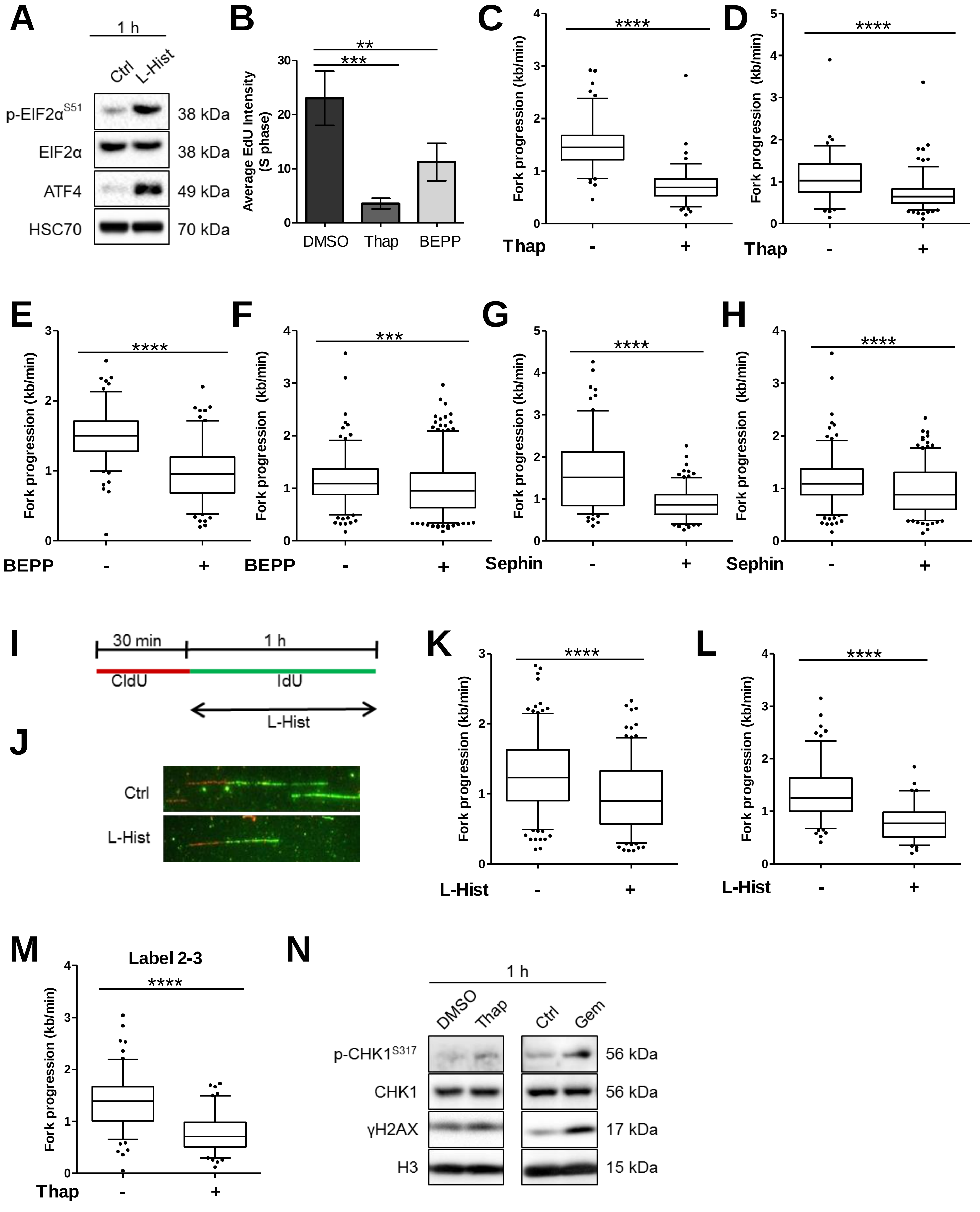

### Supplementary Figure S2

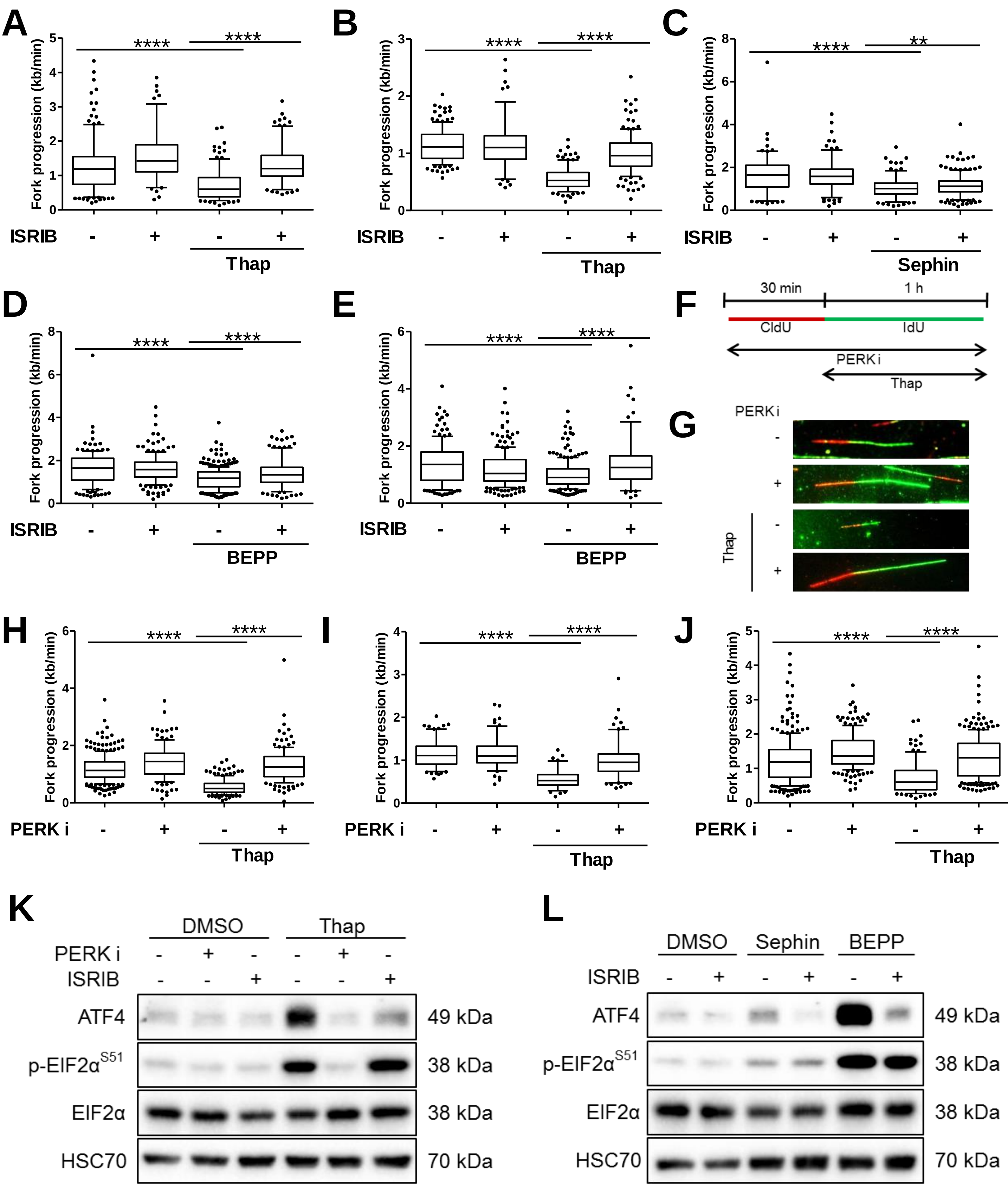

### Supplementary Figure S3

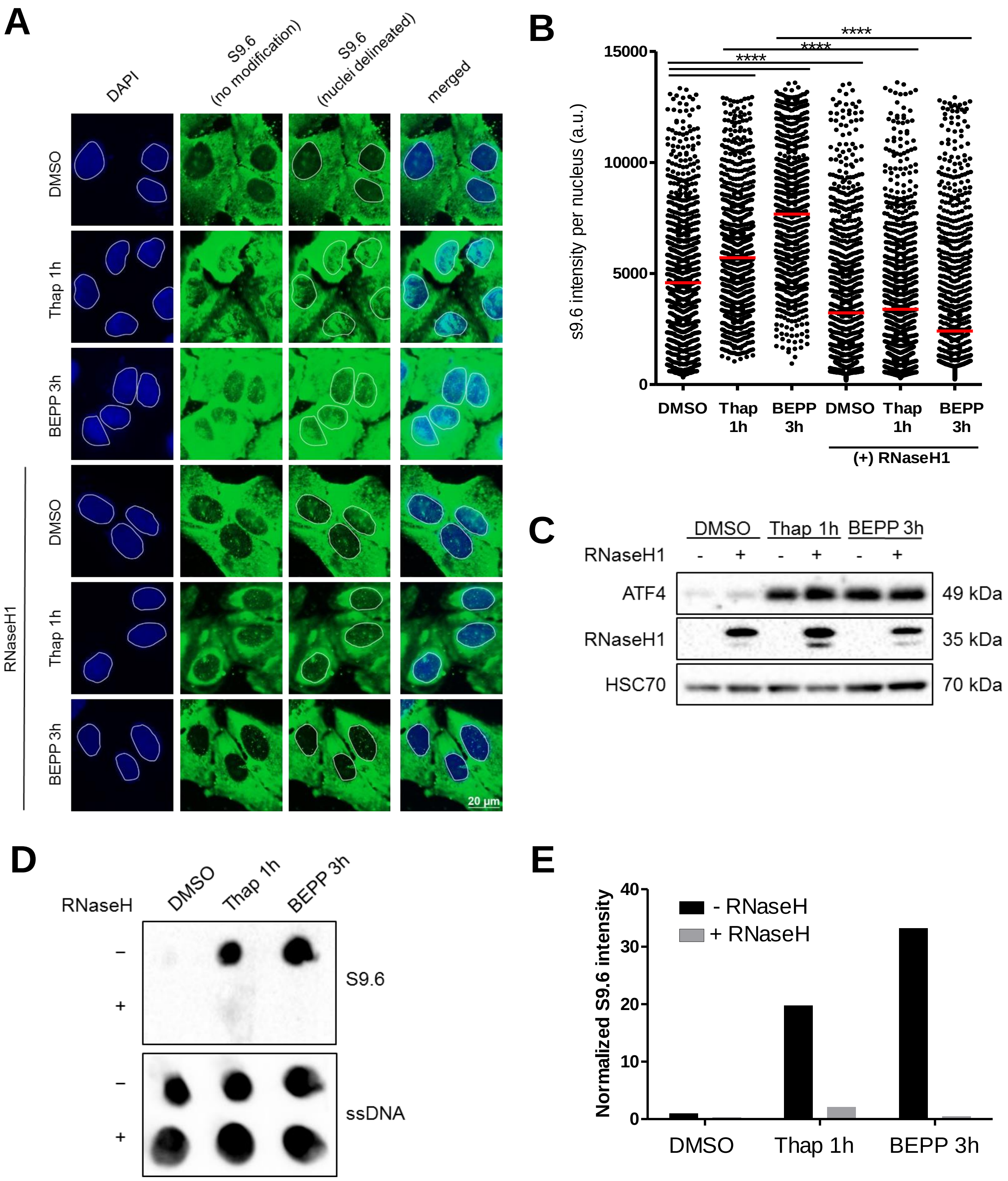

### Supplementary Figure S4

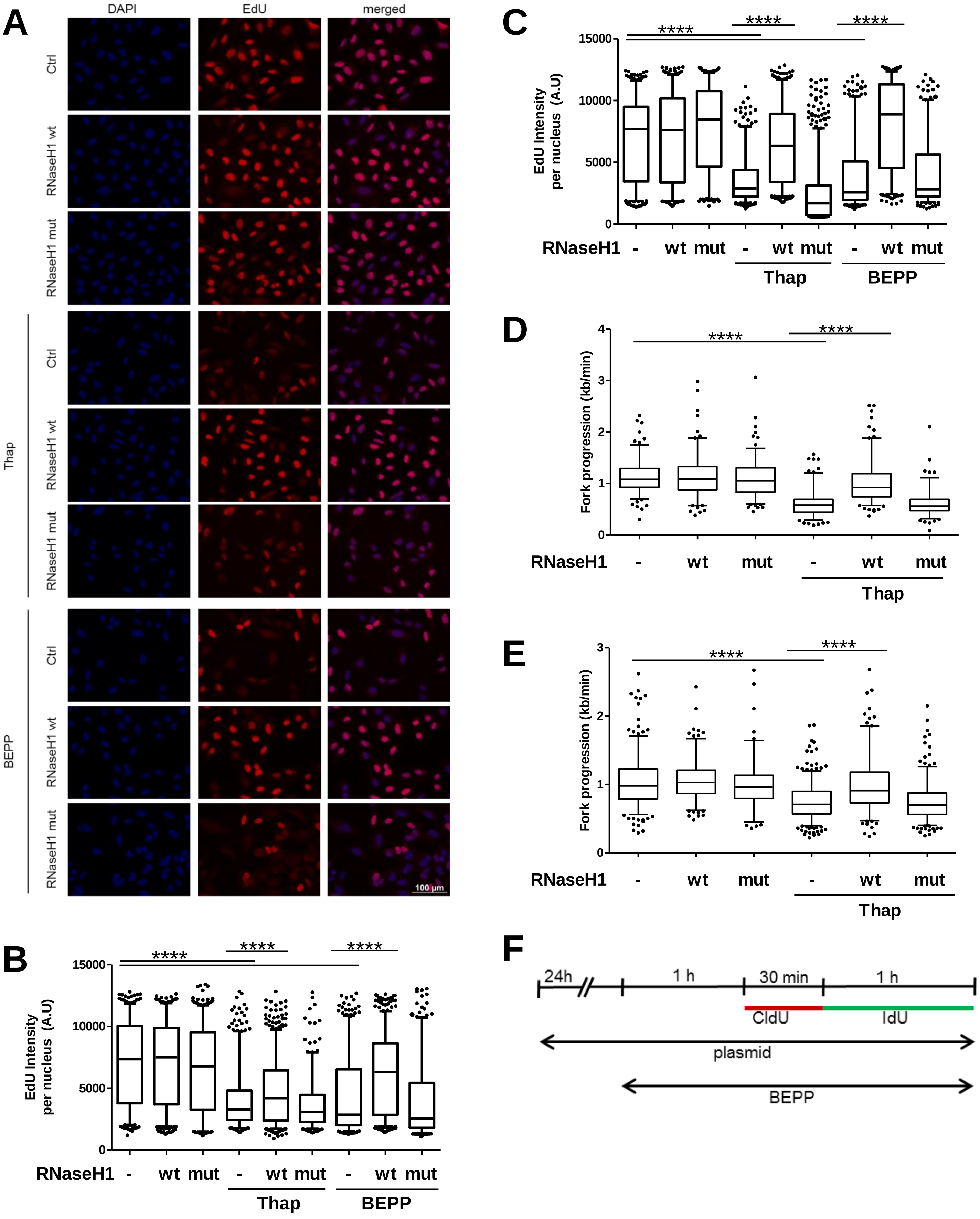

### Supplementary Figure S4 (continued)

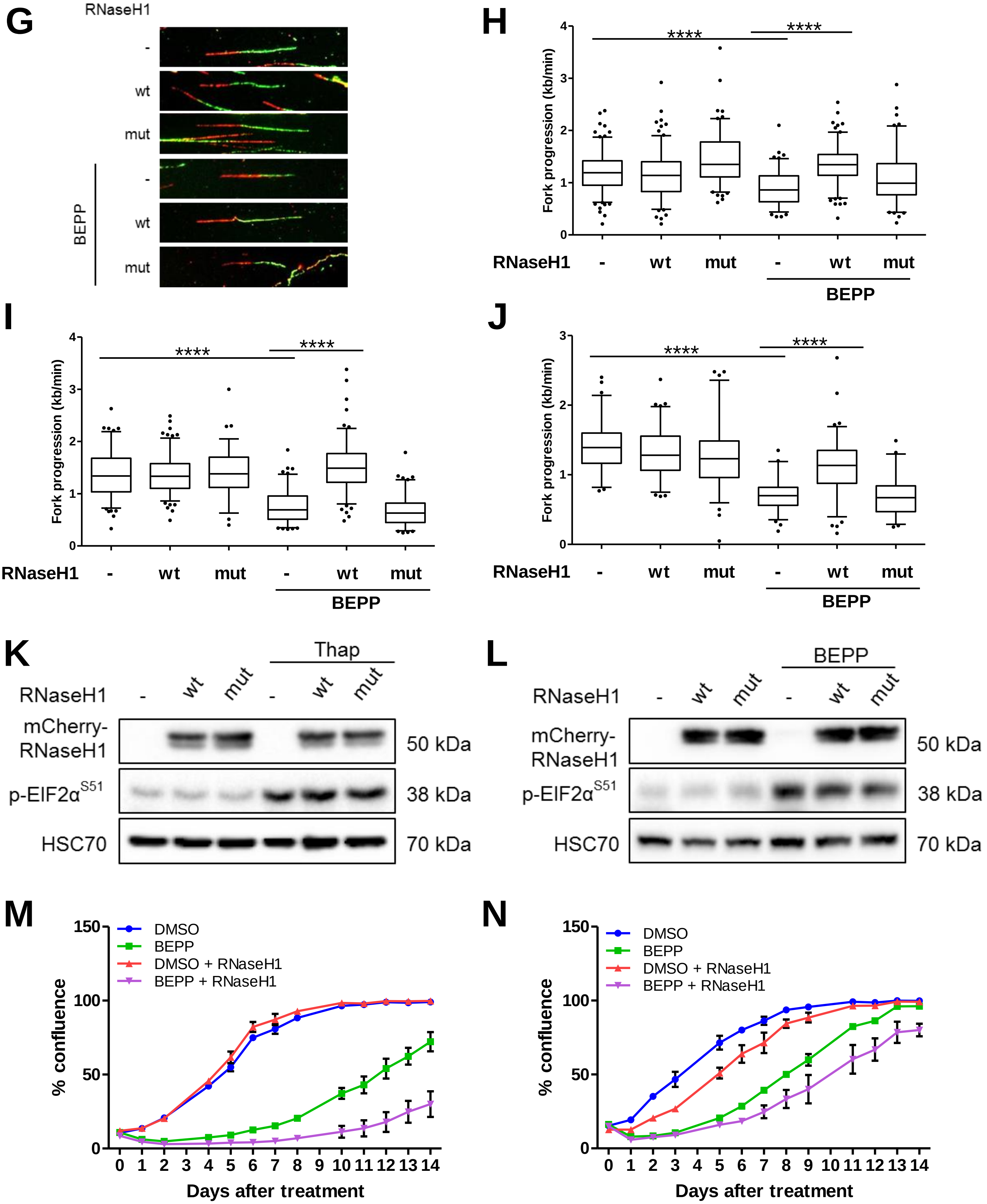

### Supplementary Figure S5

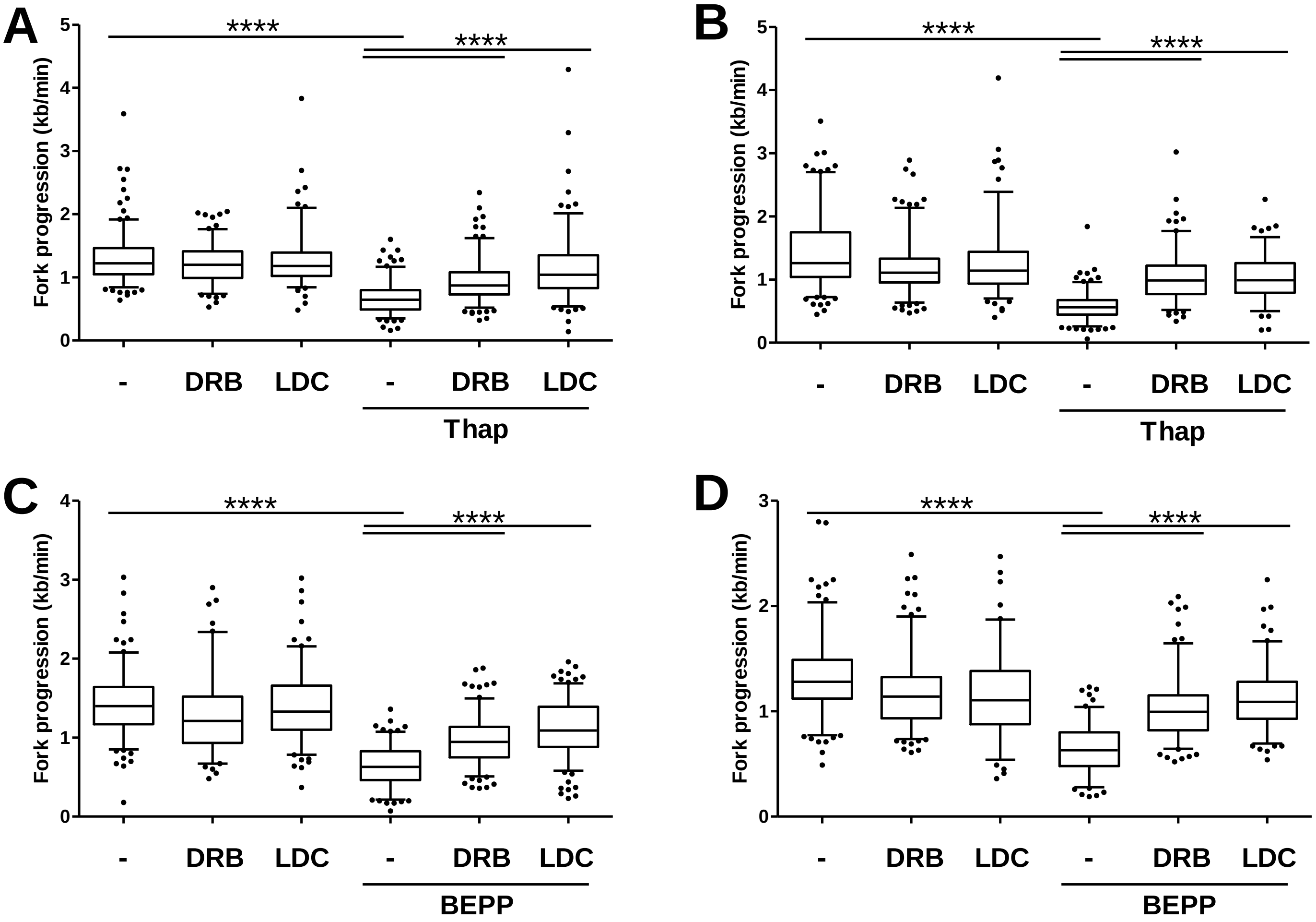

### Supplementary Figure S6

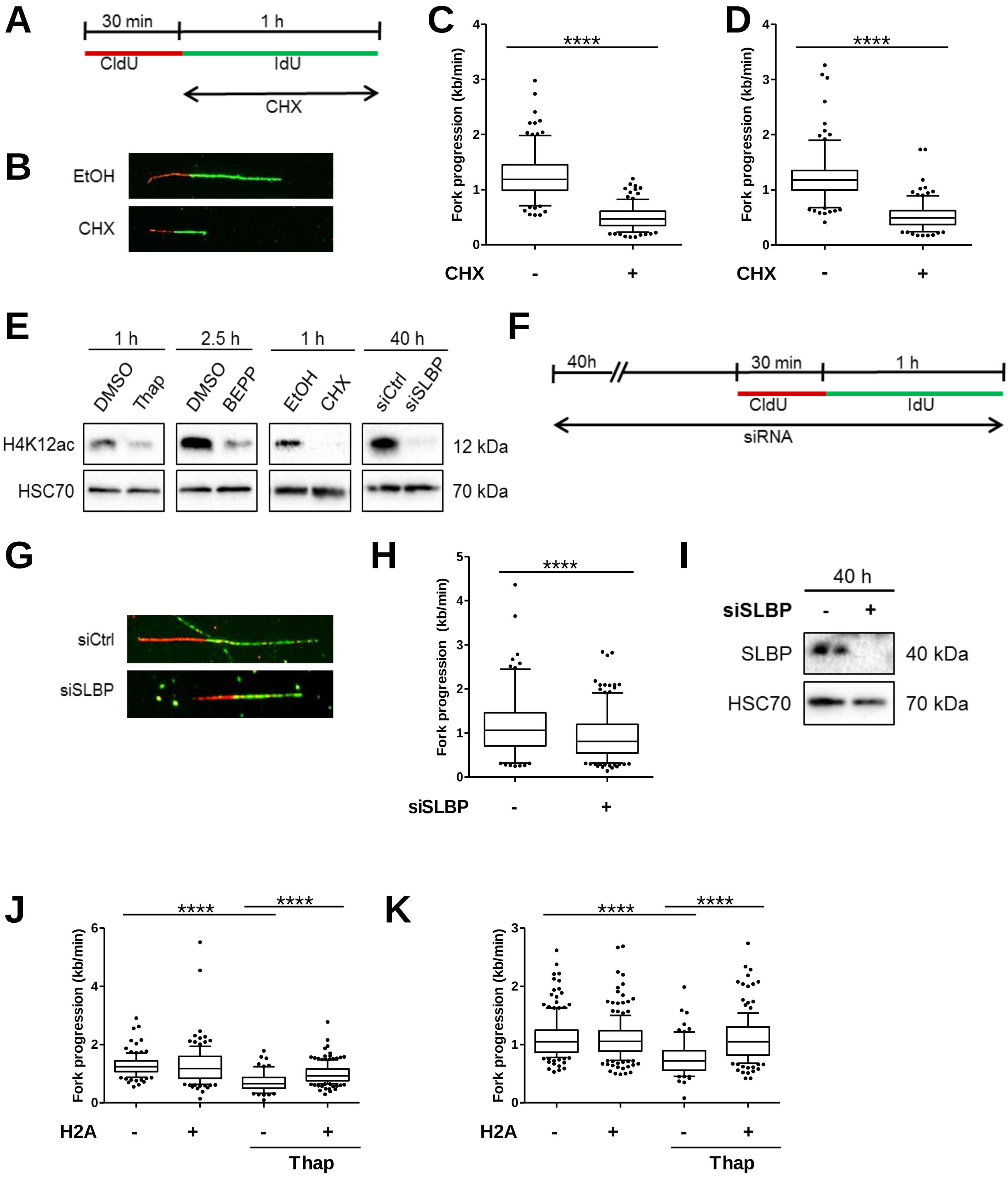

### Supplementary Figure S6 (continued)

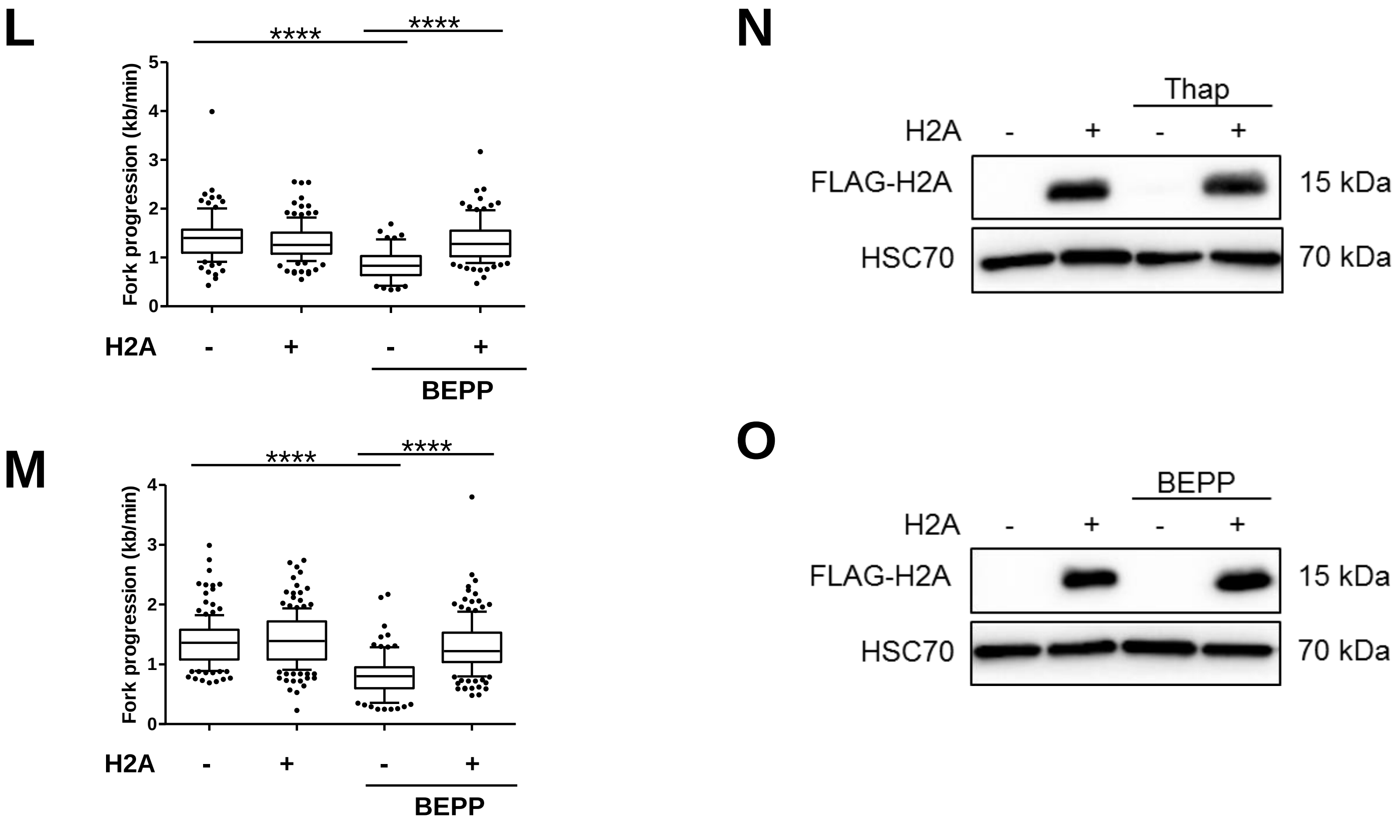

### Supplementary Figure S7

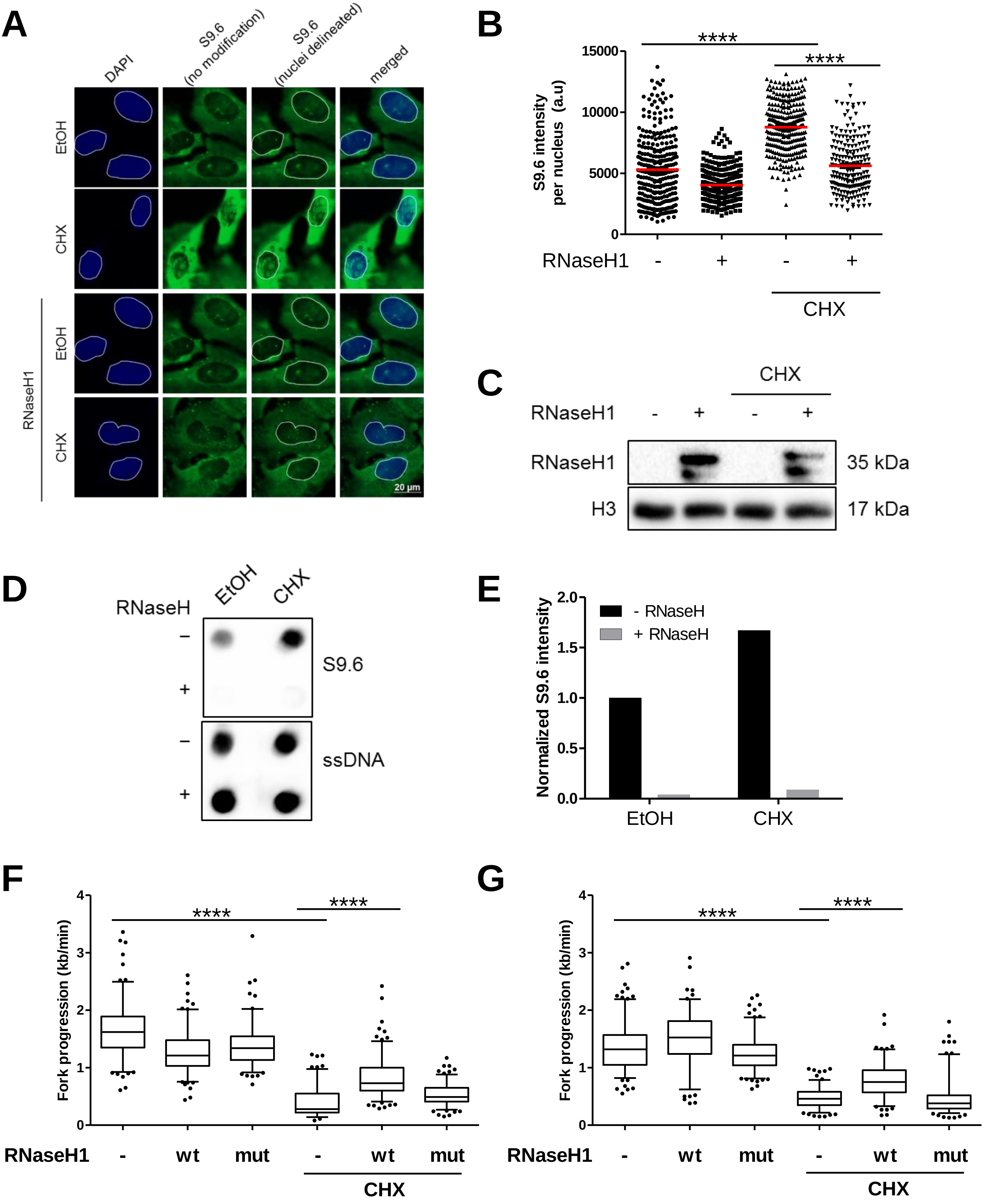

### Supplementary Figure S7 (continued)

H

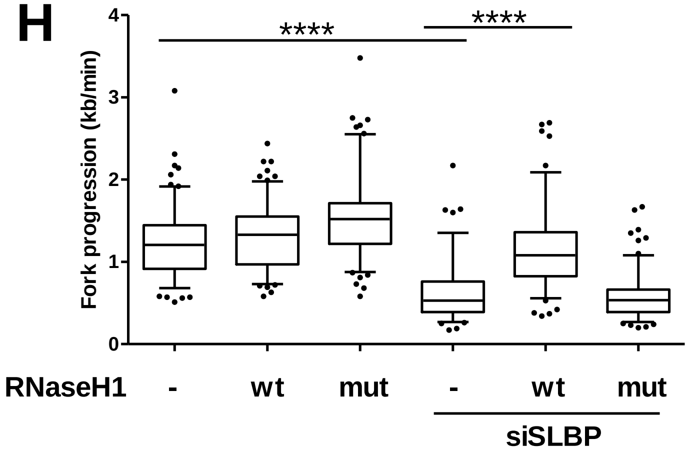


I

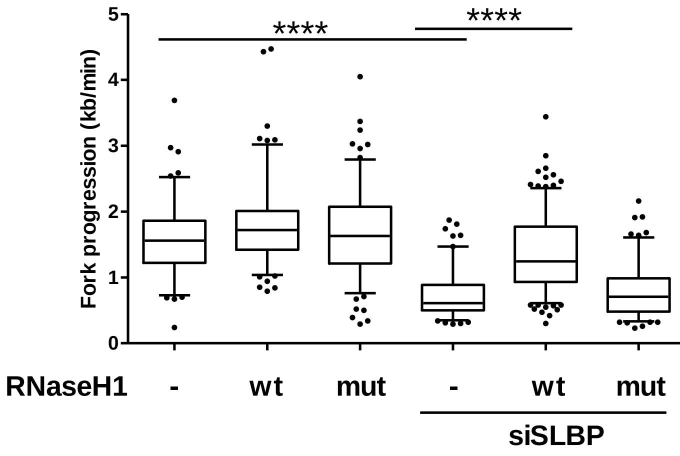


J

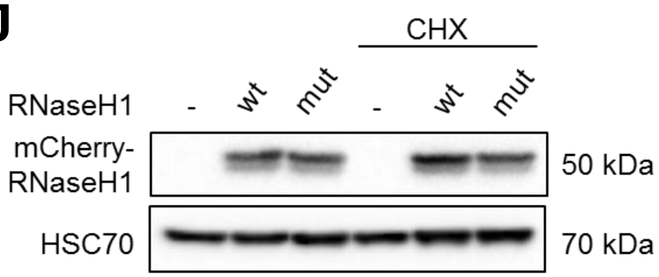


K

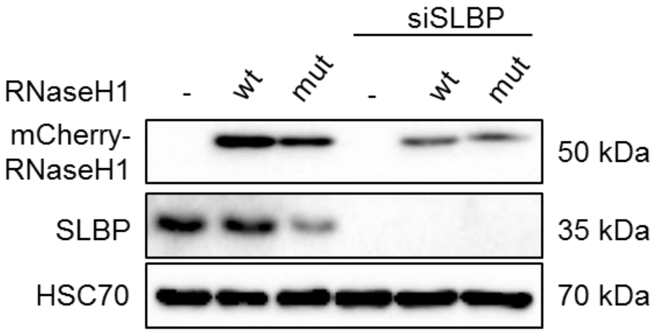
