## Supplementary figure legends for "The integrated stress response induces R-loops and hinders replication fork progression"

Running title: ISR antagonizes DNA replication

Keywords: Integrated stress response, PKR, PERK, GCN2, eIF2alpha, Thapsigargin, BEPP, ISRIB, R-loops, histones, DNA replication, DNA fiber assays

Declaration of interests: The authors declare no conflict of interests.

### SUPPLEMENTARY FIGURE LEGENDS

**SUPP FIGURE 1: DNA replication is compromised shortly after ISR induction.** *Related to Fig. 1*.

**(A)** Immunoblot analysis of cells treated with control or L-Hist (4 mM ,1 h) to confirm ISR induction (eIF2alpha phosphorylation status and ATF4 levels). HSC70 as loading control.

**(B)** Average EdU intensity of cells in S phase displayed as mean ± SD of the second independent experiment conducted in technical triplicates. See Fig. 1D.

**(C−H)** DNA fork progression (kb/min) of Thap (**C,D**), BEPP (**E,F**) or Sephin (**G,H**) – treated cells as described in Fig. 1E or 1H. Fork progression calculated using IdU track and displayed as box plots with 5-95 percentile whiskers. Replicate experiments for Fig. 1G,K,L respectively.

**(I)** U2OS cells were incubated with 5'-chloro-2'-deoxy-uridine (25 µM CldU, 30 min) followed by 5-iodo-2'-deoxyuridine (250 µM IdU, 60 min) in the presence of 4 mM L-Hist prior to harvesting for DNA fiber analysis.

**(J)** Representative images of IdU (green) and CldU (red) track lengths of L-Hist−treated cells.

**(K/L)** Box plot (5-95 percentile) of DNA fork progression (kb/min) of IdU label of cells treated with L-Hist. Cells were treated and labeled as described in (**I**). Two biological replicates of 3 shown.

**(M)** Fork progression as determined using the track length of label 2-3 of the 7-label fiber assay. Box plot with 5-95 percentile whiskers of the second biological repeat of Fig. 1P shown.

**(N)** Cells were treated with Thap (4 µM) or Gem (500 nM) for 1 h and then harvested for western blot analysis. Chk1 phosphorylation and gamma H2AX induction marks replicative stress (1). Total Chk1 levels and H3 were used as loading controls. Thap treatment does not induce Chk1 phosphorylation or gamma H2AX to a similar extent as Gemcitabine treatment, a widely used replicative stress inducer.

**SUPP FIGURE 2: Pharmacological antagonists of ISR partially rescue DNA replication.** *Related to Fig. 2.*

**(A/B)** Biological repeats of the experiment described in Fig. 2A. Box plot (5-95 percentile whiskers) showing fork progression rate (kb/min) of ISRIB/Thap–treated cells, calculated using IdU length. Corresponding to Fig. 2F**.**

**(C−E)** Box plot (5-95 percentile whiskers) showing DNA fork progression rate (kb/min) measured using IdU length of ISRIB/Sephin (**C**) or ISRIB/BEPP (**D,E**) –treated cells as described in Fig. 2B. Replicates to Fig. 2G,H**.**

(**F**) U2OS cells were treated with 0.5 μM PERK i in the presence of CldU (30 min) followed by IdU (60 min) with both PERK i and 4 μM Thap as indicated prior to DNA fiber analysis.

**(G)** DNA fiber tracks (representative) of cells in (**I**) visualized with immunostaining of CldU (red) and IdU (green).

**(H−J)** Fork progression rate (kb/min) of PERK i/Thap –treated cells of the experiment described in **(F)**. IdU label was used to calculate fork rate and displayed as box plots (5-95 percentile whiskers).

**(K)** Western blot analysis of PERK i or ISRIB–treated cells with or without Thap confirming activation and inhibition of ISR in the context of PERK stimulation. HSC70 as loading control.

**(L)** Expression of ATF4 and eIF2alpha phosphorylation status as measured *via* immunoblot analysis to ensure ISR activation and/or inhibition. HSC70 was visualized to ensure equal loading.

**SUPP FIGURE 3: Stimulation of the ISR induces R-loops.** *Related to Fig. 3.*

**(A)** Representative images of cells treated as described in Fig. 3A visualized using DAPI (nuclei) or Alexa-Fluor488 (S9.6) staining. S9.6 antibody was used to detect R-loops as described. Note that the antibody also gives rise to a fluorescence signal in the cytoplasm, in agreement with previous reports (2, 3). Unlike the nuclear signal, the cytoplasmic fluorescence was not removed by RNaseH1 and thereby confirmed to be non-specific. DAPI was used to determine the nuclei for quantification of S9.6 staining as indicated by the white outlines. Of note, the nuclear S9.6 signal was removed by RNaseH1, indicating that it truly reflects DNA:RNA hybrids. Scale bar: 20 µm.

**(B)** Biological replicate of the experiment described in Fig. 3A showing S9.6 intensity per nucleus of cells (in arbitrary units) after quantification. Red line represents mean nuclear S9.6 staining.

**(C)** Western blot analysis confirming RNaseH1 overexpression and ISR activation (ATF4 level). HSC70 as loading control.

**(D)** Replicate of the dot blot analysis for S9.6 detection to quantify R-loops and ssDNA as internal sample loading control. Samples were subjected to RNaseH treatment to confirm the specificity of the signal. See Fig. 3B***.***

**(E)** Quantification of the S9.6 signal detected in (**D**) normalized against the ssDNA and against the DMSO (without RNaseH) control. See Fig. 3C**.**

**SUPP FIGURE 4: Removal of R-loops re-establishes DNA replication upon induction of ISR but compromises survival of stressed cells.** *Related to Fig. 4.*

**(A)** Cells were treated as described in Fig. 4A. Representative images of DAPI (blue) and EdU (red) signals of one independent experiment out of 3 are shown. Scale bar: 100 µm

**(B/C)** Biological replicates showing EdU intensity per nucleus of Thap or BEPP−treated cells in the presence/absence of wildtype or mutant RNaseH1. Box plots with 5-95 percentile whiskers shown. Corresponding to Fig. 4A.

**(D/E)** DNA fork progression (kb/min) measured using IdU track length of Thap–treated cells with/without RNaseH1 overexpression plasmids as described in Fig. 4B. Results displayed as box plots with 5-95 percentile whiskers. Replicates to Fig. 4D.

**(F)** Transfection of cells with control or RNaseH1 plasmids (wt or mut) were conducted as described. Cells were then treated with 10 µM BEPP for 1 h and then incubated in CldU (25 µM, 30 min) and IdU (250 µM, 60 min) in the presence of BEPP prior to analysis.

**(G)** Labeled DNA fibers visualized *via* immunostaining of CldU (red) and IdU (green) of BEPP–treated cells overexpressing the respective plasmids described in (**F**).

**(H−J)** Box plots (5-95 percentile whiskers) showing the DNA fork progression (kb/min) of cells overexpressing RNaseH1 wt/mut and treated with BEPP as described. IdU label was used to measure fork progression.

**(K/L)** Activation of ISR and RNaseH1 overexpression confirmed with immunoblot analysis of phosphorylated eIF2alpha and mCherry respectively with HSC70 as loading control.

**(M/N)** Biological replicates of the proliferation assay described in Fig. 4E of S phase cells treated with 30 µM BEPP for 6 h with/without RNaseH1 overexpression.

**SUPP FIGURE 5: Ongoing transcription is required for compromising DNA replication by the ISR.** *Related to Fig. 5.*

**(A−D)** IdU fork progression in kb/min of cells treated with CDK9i and Thap (**A−B**) or BEPP (**C−D**) as in Fig. 5A,B displayed as box plots (5-95 percentile whiskers). Biological replicates to Fig. 5E,F respectively.

**SUPP FIGURE 6: ISR activation blocks the synthesis of histones required for DNA replication.** *Related to Fig. 6.*

**(A)** U2OS cells were labeled with CldU (25 µM, 30 min) followed by IdU (250 µM, 60 min) in the presence of cycloheximide (CHX, 50 µg/ml) and then harvested.

**(B)** Representative DNA fibers of cells treated with CHX or solvent control (EtOH) visualized with immunostaining of CldU (red) and IdU (green).

**(C/D)** Box plot (5-95 percentile whiskers) showing fork progression (kb/min) calculated using IdU track length of CHX–treated cells in **(A).** Two biological replicates of 3 shown.

**(E)** Soluble proteins were extracted as described in Fig. 6A from Thap (4 µM), BEPP (10 µM), CHX (50 µg/ml) –treated cells or cells transfected with siRNA against SLBP (100 nM). Immunoblot analysis of soluble histone-4 lysine-12 acetylation (H4K12ac) was used as a mark for newly synthesized histones. HSC70 was used as loading control.

**(F)** U2OS cells were transfected with control or siRNA against SLBP (100 nM) for 40 hours prior to incubation with CldU (25 µM, 30 min) and IdU (250 µM, 60 min). Cells were the harvested for DNA fiber analysis.

**(G)** Representative DNA fiber tracks of cells treated depleted of SLBP visualized via immunostaining of CldU (red) and IdU (green).

**(H)** IdU track length of cells in **(G)** was used to measure DNA fork progression (kb/min) as displayed as box plots (5-95 percentile whiskers). One representative experiment out of 3 shown.

**(I)** Western blot analysis of cells treated in **(F)** confirming SLBP knock down. HSC70 used as loading control.

**(J/K)** Box plots (5-95 percentile whiskers) showing DNA fork progression (kb/min) measured using the IdU tracks of cells treated with Thap and transfected with H2A plasmid as described (Fig. 6B). Replicates to Fig. 6F.

**(L/M)** DNA fork progression (kb/min) of cells treated as described in Fig. 6C and displayed as box plots (5-95 percentile whiskers). IdU label was used to calculate fork progression. Replicates to Fig. 6G.

**(N/O)** Immunoblot analysis of cells confirming H2A overexpression (Flag) in cells treated with Thap **(N)** or BEPP **(O).** HSC70 was used as loading control.

**SUPP FIGURE 7: Inhibition of histone synthesis induces R-loops which impairs DNA replication.** *Related to Fig. 7.*

**(A)** S phase cells overexpressing control or RNaseH1 plasmids were treated with CHX (50 µg/ml) for 1 h and then harvest for S9.6 immunofluorescence analysis as described. Representative images of cells as visualized using DAPI (nuclei) or Alexa-Fluor 488 (S9.6) staining to detect R-loops. DAPI was used to determine regions of interests within the nuclei for quantification as indicated by the white outlines. Scale bar: 20 µm.

**(B)** Biological replicate of S9.6 immunofluorescence staining of CHX−treated cells quantified and plotted as scatted plots. Red line represents mean S9.6 intensity per nucleus. Corresponding to Fig. 7A**.**

**(C)** Western blot analysis confirming RNaseH1 overexpression in cells describe in Fig. 7A and in (**B**). Total H3 used as loading control.

**(D/E)** Biological replicate of the dot blot analysis in Fig. 7B,C. S9.6 signal intensity in (**D**) was quantified and normalized to the internal loading control (ssDNA), then to the EtOH control (without RNaseH) and plotted as bar charts (**E**).

**(F/G)** Box plots with 5-95 percentile whiskers displaying the DNA fork progression of cells treated/transfected as in Fig. 7D. IdU track length was used to calculate the fork progression of CHX−treated cells with RNaseH1 overexpression. Replicates to Fig. 7H.

**(H/I)** Cells transfected with siRNA/plasmid as described in Fig. 7E were harvested for DNA fiber analysis upon labeling with CldU and IdU. DNA fork progression of the IdU label (kb/min) of the additional 2 independent experiments corresponding to Fig. 7I shown as box plots (5-95 percentile whiskers).

**(J)** Immunoblot analysis of mCherry confirming overexpression of RNaseH1 in cells described in Fig. 7D.

**(K)** RNaseH1 overexpression and SLBP knockdown were confirmed with immunoblot analysis to mCherry and SLBP respectively. HSC70 as loading control.

**REFERENCES**


1. Dobbelstein M, Sorensen CS. Exploiting replicative stress to treat cancer. Nat Rev Drug Discov. 2015;14(6):405-23.

2. Salas-Armenteros I, Perez-Calero C, Bayona-Feliu A, Tumini E, Luna R, Aguilera A. Human THO-Sin3A interaction reveals new mechanisms to prevent R-loops that cause genome instability. EMBO J. 2017;36(23):3532-47.

3. Schwab RA, Nieminuszczy J, Shah F, Langton J, Lopez Martinez D, Liang CC, et al. The Fanconi Anemia Pathway Maintains Genome Stability by Coordinating Replication and Transcription. Mol Cell. 2015;60(3):351-61.
